# Takeoff dynamics are stereotyped across jumping spiders

**DOI:** 10.64898/2026.08.19.745647

**Authors:** Erin E. Brandt, Alasdair D. Hastewell, Lin Yan, Kaci Rose Goldberg, Jacob S. Harrison, Lizbeth Badillo Aguilar, Damian O. Elias, M. Saad Bhamla, Jasmine Nirody

## Abstract

Jumping is a challenging locomotive mode, requiring rapid force generation and precise co-ordination of multiple limbs. Many animals meet this challenge using elastic mechanisms that store and rapidly release energy. Jumping spiders (Salticidae), however, rely on a semi-hydraulic system that constrains how their legs can generate propulsion. Much about how these spiders reliably generate jumps within these mechanical constraints remains unknown. Here, we analyze 46 individuals collected from the Peruvian Amazon, spanning 14 genera and significant morphological diversity, and show that this physically constrained system is coupled to a remarkably stereotyped coordination strategy. Whole-body kinematics and novel graph-based analyses of inter-limb coordination reveal a stereotyped two-stage takeoff sequence: a “swing” driven by extension of the fourth legs, followed by a rapid “fling” by the third legs that generates propulsion for takeoff. We further demonstrate that this pattern is preserved beyond Amazonian species, persisting in salticids from North America and Australia. Our results suggest that physical and biomechanical constraints may canalize locomotor evolution toward shared dynamical solutions.

## Introduction

Jumping is a demanding mode of locomotion. Successful takeoff requires the rapid generation of large forces while coordinating multiple limbs with an extremely high level of precision [3]. Across the animal kingdom, evolution has repeatedly met this challenge through various elastic mechanisms that store energy and release it rapidly, enabling movements that exceed the limits of muscle alone [5, 9, 8, 42, 27, 15]. Yet not all animal morphologies allow for spring-driven power amplification mechanisms. Understanding how successful locomotive strategies evolve under such physical constraints remains a fundamental challenge in biomechanics and evolutionary biology.

Jumping spiders are a striking example of this: rather than relying primarily on elastic recoil, they generate propulsion using a combination of direct muscular actuation and hydraulically-driven leg extension to power takeoff [30, 16, 46, 6]. This semi-hydraulic system imposes physical constraints on the speed and force with which the legs can be actuated, likely limiting the mechanical strategies available for generating rapid takeoff [45, 25, 38, 6].

Despite these constraints, salticids have radiated into approximately 7,000 species spanning extraordinary diversity in body size, morphology, habitat, and ecology – all while relying extensively on jumping [22]. These spiders routinely jump to traverse discontinuous vegetation, capture prey, evade predators, and navigate complex environments [11, 2, 47, 10]. In contrast to many other pervasive biomechanical systems, their success appears to be due not to maximizing any single performance metric but rather to their ability to execute jumps reliably within a constrained mechanical landscape [25, 6].

But how do jumping spiders consistently generate effective jumps given these limitations? The paradoxical nature of their success raises the broader question: When physical constraints limit the mechanical solutions available for a behavior, does evolution tend to discover different specialized coordination strategies, or does it converge on a common motor solution that remains effective despite morphological and ecological diversification? In the case of salticids, do different species achieve this through distinct lineage-specific strategies, or is there a common biomechanical paradigm that enables robust and reliable performance across variation in morphology, size, and ecology?

Here, we take advantage of recent advances in high-speed videography and deep-learning pose estimation techniques [24, 33] to analyze body motion and mechanisms of force production during takeoff in Amazonian jumping spiders spanning 14 genera and remarkably broad morphological diversity. We develop a novel graph-based framework for analyzing limbed locomotion that explicitly incorporates the topology of the spider body plan. By treating limbs and joints as nodes within a dynamical structured network, this sensitive approach captures patterns of intra-body movement and inter-limb coordination that are difficult to resolve with conventional kinematic metrics alone.

Our analyses uncover that jumping spiders across substantial diversity in phylogeny, morphology, and ecology utilize strongly conserved intra-body coordination patterns during takeoff. These results suggest that successful spider jumps likely rely on a shared motor template that operates robustly within the physical constraints imposed by their semi-hydraulic actuation. A candidate mechanism is a coordinated “swing-and-fling”, in which a sequence of leg movements first generate body opening and loading and then rapidly extend to drive propulsion. This work presents a unifying framework that can reveal conserved principles of movement across both biological, robotic, and other mechanical systems, using salticid jumps as an example where morphology can shape locomotor evolution by constraining the diversity of viable coordination strategies.

## Results

### Amazonian jumping spiders span substantial morphological diversity

We analyzed 46 individuals representing 14 genera of Amazonian jumping spiders (Figure 1). Many of these individuals represent undescribed species within a described genus and therefore were identified to genus. The animals analyzed here encompass striking diversity in body size, leg proportions, and coloration. Body lengths, measured from the anterior tip of cephalothorax to the posterior tip of the abdomen, ranged from 4.6–9.5 mm (mean 6.8 ± 1.2 mm), while leg lengths varied nearly threefold (3.6–10.7 mm, with mean 6.5 ± 1.5 mm). Overall, females in our sample were larger than males.

**Figure 1:**
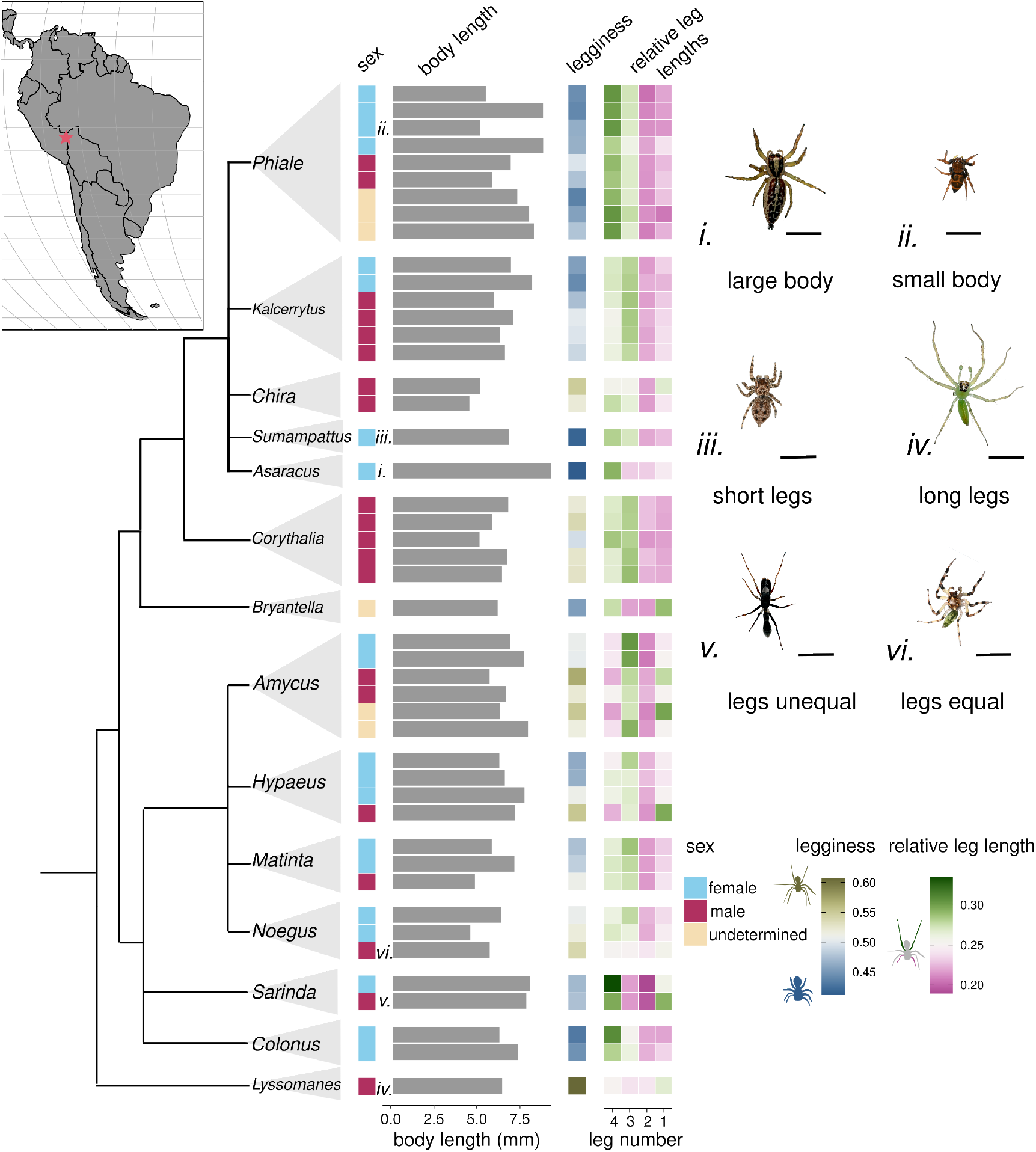
Phylogenetic and morphological spread of sampled Amazonian salticids. Map in upper left indicates location of sampling: Finca las Piedras, Peru. Phylogeny of sampled genera (left, n = 14, tree branch lengths not to scale) alongside quantitative summaries of various morphological parameters (center) and representative phenotypes (right). For each individual, the following metrics are shown: sex (male, female, undetermined), body length (mm), legginess, and relative leg lengths. Body length is measured from the anterior margin of the cephalothorax (head) to the posterior tip of the abdomen. Legginess is defined as the ratio of body length to mean leg length. Relative leg length *R*_*i*_ for each leg pair *L*_*i*_, *i* = 1, 2, 3, 4 is computed as follows: 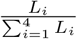; equally-sized leg pairs correspond to *R*_*i*_ = 0.25 for all *i*. See Supplementary Figure S1 for more details on these metrics. Heatmaps summarize variation in legginess and the distribution of relative leg lengths across leg pairs across sampled genera. Representative images (i–iv) illustrate extremes and qualitative categories captured by these metrics (scale bar = 5 mm in all images). Corresponding genera are noted along the phylogeny. See Supplementary Figure S2 for images of all photographed individuals.

To quantify morphological variation independently of overall size, we further defined two metrics of relative leg length: *legginess*, the ratio of average leg length to body length, and *leg ratio* (see Supplementary Figure S1), the relative contribution of each leg pair to total leg length. The average legginess was 1.04 ± 0.17, with a range of 0.64 - 1.42. Leg ratio is the relative length of legs relative to the others. A value of 0.25 for each leg indicates all legs of equal length. Values below 0.25 represent a relatively shorter leg, and values above 0.25 represent a relatively longer leg. The average leg ratio was 0.25 ± 0.3 SD, with a range of 0.19 - 0.33. These metrics reveal broad variation in overall body plan, providing an opportunity to test whether jumping dynamics diversify alongside morphology.

### Jump kinematics are tightly conserved across salticid diversity

We recorded spiders jumping from the edge of a platform using high speed cameras at 1,000 fps (Figure 2A). For each video, 25 keypoints corresponding to anatomical landmarks were tracked (5 along each leg on the right side and 5 on the body). We estimated whole-body kinematics using the mean position of the body landmarks as a proxy for the center of mass; to account for jitter in landmark localization, we used a custom smoothing spline approach (details in Supplementary Information). Despite the substantial morphological diversity of the spiders, velocity and acceleration profiles exhibited a strikingly similar temporal structure across spiders (insets, Figure 2B).

**Figure 2:**
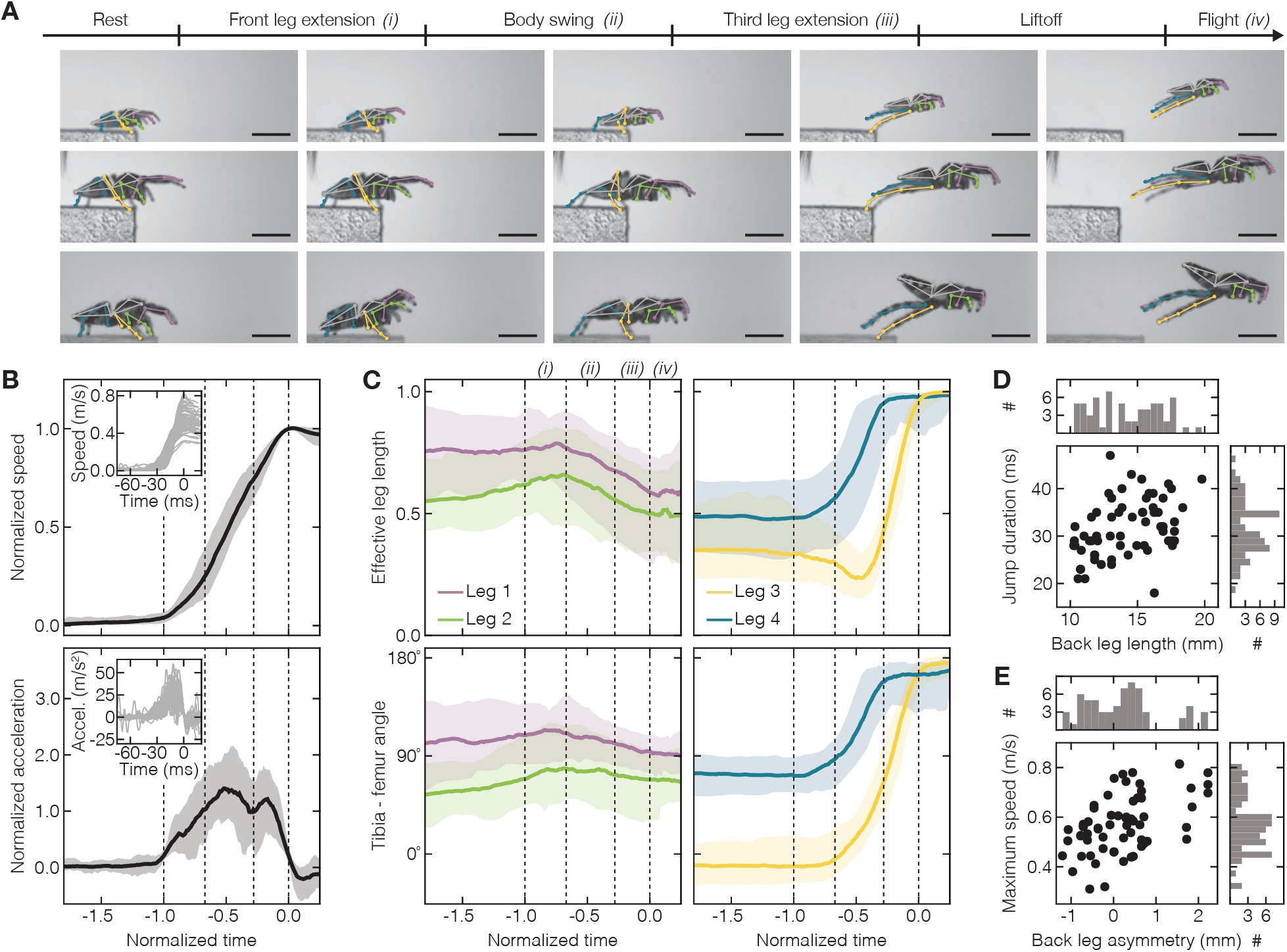
Jump kinematics converge across diverse body plans. **A** Representative jumping sequence showing the conserved major phases of takeoff of a representative salticid jump. Scale bar 5mm. See Supplementary Figure S3 for jumping sequence of all 66 videos. **B** Whole-body speed (top) and acceleration (bottom) during jumping. Individual jumps show highly similar kinematic profiles despite variation among spiders. Shaded regions show 5th to 95th percentile range with the median as solid line. Nondimensionalization by jump duration and maximum speed reveals a strongly conserved temporal profile across individuals; insets show data before nondimensionalization. The characteristic double peak in acceleration suggests two distinct phases of takeoff. **C** Dynamics of effective leg length (top) and tibia-femur angle (bottom) for all four leg pairs. The first and second legs show greater variability before takeoff but with a characteristic peak in extension right before fourth leg extension *(i)*. The third and fourth legs undergo the most dramatic changes during takeoff and exhibit a stereotyped temporal sequence, with fourth-leg extension *(ii)* preceding rapid extension of the third legs *(iii)*. See Supplementary Figure S4 for individual traces. **D** Maximum jump speed scales with asymmetry between the third and fourth legs’ length. **E** Jump duration scales with the combined length of the third and fourth legs’ length. Together, these results indicate that while morphology sets the characteristic timescales of jumping, the temporal organization of takeoff is highly conserved.

We found that particular features of jump kinematics were tightly linked to lengths of the spiders’ third and fourth leg pairs (*ℓ*_3_ and *ℓ*_4_, respectively): the jump duration depends on the combined length of the third and fourth legs *ℓ*_3_ + *ℓ*_4_ (Figure 2D) and the maximum speed depends on asymmetry between these two leg pairs *ℓ*_3_ −*ℓ*_4_ (Figure 2E). To explore the similarity of the temporal organization of jumps separate from these characteristic scaling relationships, we nondimensionalized kinematic measurements using jump duration and maximum jump speed (see Materials and Methods for more details). Following this normalization, kinematic trajectories from individuals spanning the full range of body sizes and morphologies collapsed onto a highly stereotyped profile (Figure 2B,C). These results suggest that, while leg morphology may regulate the timescale and peak speed of takeoff, the force generation mechanism underlying jumps is likely conserved across spider phylogeny and morphology.

The acceleration profiles further revealed a reproducible double peak, suggesting that takeoff is composed of at least two temporally distinct phases (Figure 2A,B). This raised the possibility that the conserved whole-body dynamics arise from a conserved sequence of limb movements.

### A two-stage “swing-and-fling” program coordinates takeoff

To more deeply analyze the mechanism driving the double acceleration peak in our kinematic analyses, we examined the dynamics of individual limbs during takeoff. We characterized the *effective leg length*, a recently introduced measurement of leg extension, given by the ratio of the end to end distance and total leg length and the angle between the tibia and the femur. Consistent with previous observations, we find that the third and fourth legs underwent the largest changes in effective leg length during jumping and are highly stereotyped in their dynamics (Figure 2C) [6].

The movements of these two leg pairs, however, are not synchronous. Instead, effective leg length and joint angle revealed a reproducible sequence across individuals: extension of the fourth legs preceded the rapid extension of the third legs. This sequence provides a mechanical interpretation for the two peaks observed in whole-body acceleration. During the first phase of takeoff, extension of the fourth legs swings the body forward while repositioning the third legs. This is followed by rapid extension of the third legs, which generates the final propulsive impulse and launches the body from the substrate. We refer to this conserved two-stage coordination strategy as “swing-and-fling” (Figure 2A,B,C).

### Graph-based analysis reveals strongly stereotyped intra-body coordination across scales and species

We further assessed the role of whole-body and inter-limb coordination in jumping by introducing a graph-based representation of spider movement. We represented the tracked skeleton as a graph, in which the anatomical landmarks are the nodes with morphologically relevant connections, such as limbs, forming edges between them (Figure 3A).

**Figure 3:**
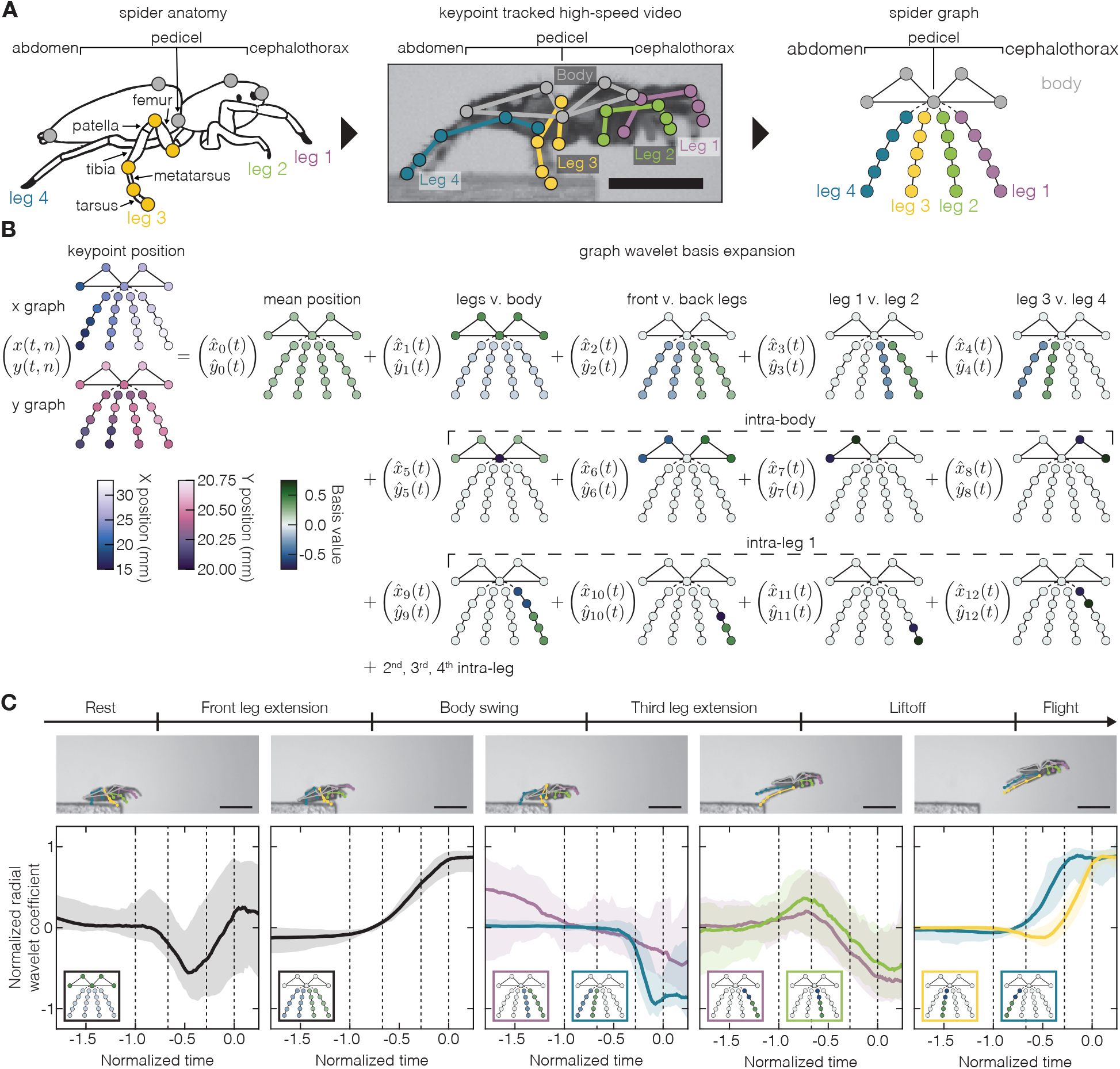
Graph-based representation reveals conserved whole-body coordination during jumping. **A** Construction of morphology-aware graph representation of spider movement from keypoint tracking. Anatomical landmarks are represented as nodes connected according to the spider body plan. **B** Spider movement is decomposed into orthogonal modes spanning progressively finer spatial scales, from whole-body motion to inter-limb and intra-limb movements. Temporal information about the inter-limb coordination is contain<u>ed in t</u>he coefficients 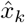 and 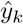. **C** Normalized rotationally invariant graph-wavelet coefficients 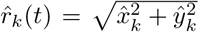 across individuals for modes describing coordination between the body and limbs (first panel), between front and back legs (second panel), between individual leg pairs (third panel), and within single legs (fourth and fifth panels). Results for the single leg modes are consistent with the previous effective leg length analysis. Mode expansions additionally reveal stereotypy between the back two legs and between the back two legs and front two legs (second and third panels). A characteristic dip between the legs and the body consistent with the “swing” phase of the jump is also observed (first panel). Dotted lines on each plot correspond to each phase of the jump takeoff, as in Figure 2. See Supplementary Figure S5 for individual traces.

We then constructed a morphology-aware sparse orthogonal basis on the nodes of this graph, motivated by Haar wavelets [23, 12, 14] (see Materials and Methods for more details). This allows us to rigorously decompose the intra-body coordination of the spiders during jumping in an interpretable way, with basis functions representing coordination at progressively finer spatial scales, from whole-body displacement to inter-limb and intra-limb motion [44, 40, 4]. For instance, the first constant basis function corresponds to the mean position of the key points, a proxy for the center of mass, while higher order bases correspond to smaller scale differences from inter-leg differences down to intra-leg and differences between individual joints within a single leg (Figure 3B).

Our wavelet analysis revealed a conserved coordination program across the whole body, with inter-limb coordination being highly stereotyped across species during takeoff. The coefficients representing the separation between the front and back legs and the distances between the back legs and front legs are strongly stereotyped over the course of the jump, with the distance between the front two legs showing more variation. These results confirm the fundamental role of the third and fourth leg pairs noted in our effective-leg-length calculations.

Furthermore, the graph analysis revealed coordination patterns that were less apparent from standard kinematic measurements. Intriguingly, the front two legs become increasingly synchronized over the course of the jump, despite showing greater variability before takeoff. This, combined with the observation that their extension peaks before the onset of the activation of the back legs, suggests a role in jump planning and preparing or stabilizing the body before propulsion. Altogether, our results indicate that swing-and-fling is not simply a sequential movement of the two back leg pairs, but part of a coordinated whole-body program.

## Discussion

Animals solve the difficult problem of jumping through a diversity of mechanical designs. Elastic catapults and latch-mediated mechanisms are common among arthropods and other small animals [5, 9, 1, 26, 7, 27, 13, 32, 19, 15]. Larger animals, which have more muscular work available to power jumps, often use direct muscular actuation [42, 43]. Each method represents a distinct evolutionary solution to the challenge of generating rapid, high-power movement. Jumping spiders, however, occupy an unusual region of this design space: they have small muscles and a semi-hydraulic locomotor system that limits opportunities for a power amplification system [18, 25, 35, 21, 6] (but see [46]). How has such a mechanically constrained system persisted in 7,000 ecologically and morphologically diverse species, all of which use jumping as their primary mode of locomotion, predator escape, and prey capture?

Here, we show that an unexpectedly conserved coordination program underlies this remarkable diversity. Across spiders varying substantially in body size and shape, phylogenetic history, and ecology (Figure 1), takeoff kinematics collapse onto highly stereotyped trajectories (Figure 2,3), with morphology impacting primarily the timescale rather than the dynamics of takeoff (Figure 2D,E). Our comprehensive and complementary analyses of body and leg kinematics and graph-based intra-body coordination patterns converge on the same conclusion: successful takeoff in jumping spiders is organized by a common temporal sequence of whole-body and inter-limb coordination. We further compared our measurements with previously published jumping data from salticids sampled across geographically disparate lineages [6, 28], the characteristic temporal structure of body speed, acceleration, and inter-limb coordination was preserved (Figure 4). This broad geographic comparison further suggests that this shared dynamical template likely extends well beyond the Amazonian assemblage studied here.

**Figure 4:**
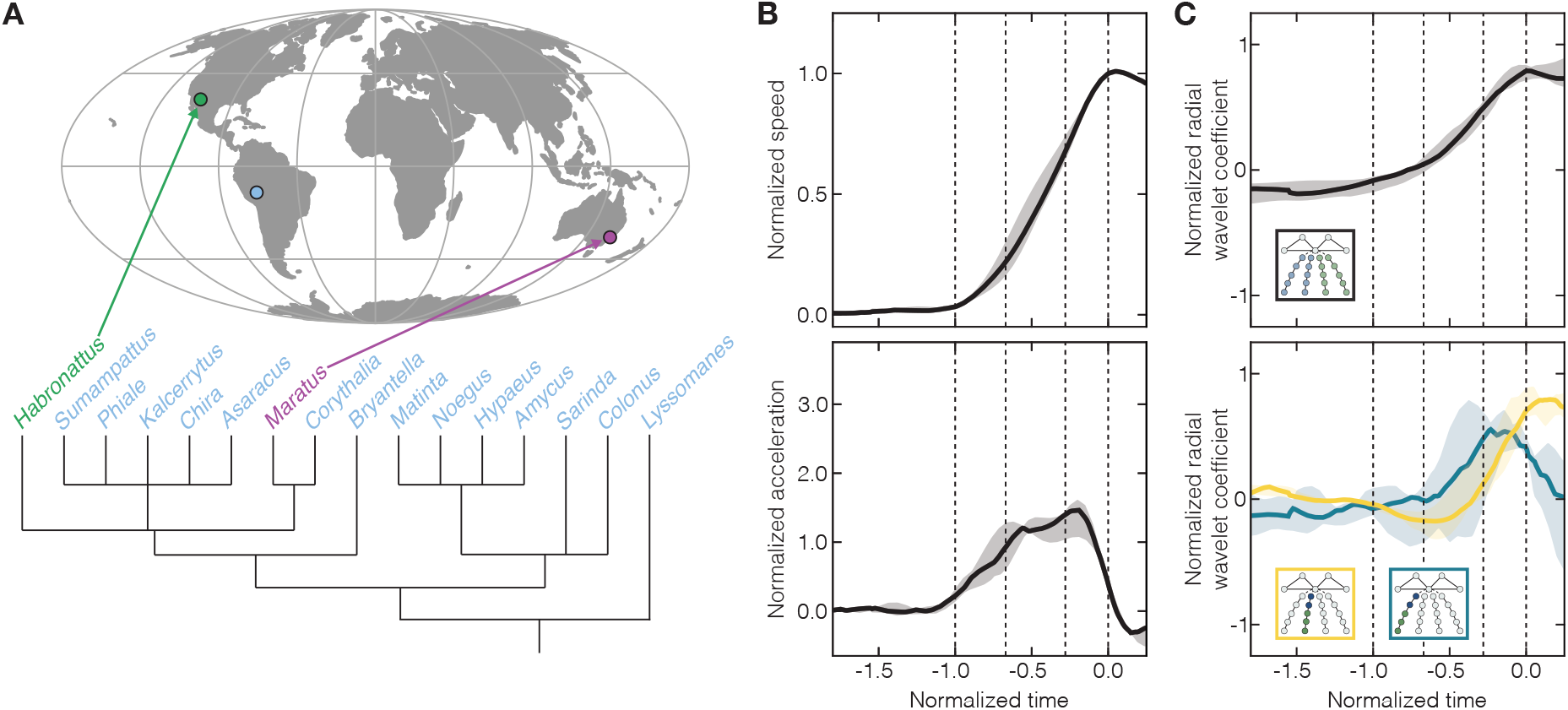
Comparison of both take-off kinematics and inter-limb coordination between the Peru genera and two additional genera – *Habronattus* from North America and *Maratus* from Australia – reveals striking similarity emphasizing that our findings span phylogenetic, morphological and also geographic distance.

Our results support a two-stage “swing-and-fling” mechanism underlying this conserved template. In this model, extension of the fourth legs first positions the body. This preparatory “swing” phase is followed by rapid extension of the third legs (“fling”), which provides the final propulsive impulse for takeoff. The reproducibility of this sequence across phylogenetically distant taxa suggests that it represents a fundamental organizational principle of salticid jumping, and could provide a testable hypothesis for how hydraulic actuation can be coordinated to produce reliable jumps in both biological and biomimetic systems [20, 39, 21, 17, 48]. Whether this sequence reflects neural control, passive mechanical coupling, or interactions between the two remains an important question for future work.

This stereotyped coordination pattern highlights the role of physical constraints imposed by the spiders’ semi-hydraulic musculoskeletal system, which fundamentally distinguishes salticids from more well-characterized elastic-powered jumping systems [18, 32]. In fleas, froghoppers, and other such specialized jumpers, elastic energy is accumulated before takeoff and released through rapid propulsion [5, 1, 26, 7, 27, 13, 32, 19, 15]. Salticids instead distribute takeoff across a sequential interaction between leg pairs, with one set of limbs repositioning the body before the second generates the final propulsive impulse. This broad conservation emphasizes how mechanical architecture of a locomotor system can substantially limit the space of viable solutions, funneling evolution toward a common motor strategy. By revealing a conserved dynamical program operating across diverse morphologies, our work highlights how physical constraints can canalize locomotor evolution and demonstrates the power of topology-aware analyses for uncovering universal principles of coordinated movement.

More broadly, our graph-based framework provides a method for quantifying coordinated movement while preserving the topology of the underlying body plan. Rather than treating individual joints or limbs independently, the representation decomposes movement into interpretable modes, from whole-body organization through inter-limb coordination to local joint movements. Because this representation is defined by anatomical connectivity, the framework provides a principled basis for comparing locomotor dynamics across diverse animals. Applying such morphology-aware approaches across taxa may help distinguish conserved motor programs from lineage-specific innovations and reveal how mechanical constraints, morphology, and control interact to shape the evolution of movement.

## Materials and Methods

### Spider collection and video recording

Data were collected from 08/04/2024 to 08/10/2024 in Finca las Piedras, Peru, as part of the Jungle Biomechanics Lab [41]. A total of 63 animals were hand collected from vegetation and building exteriors. Spiders were maintained at the research station until videos were recorded. Videos of animals jumping between two horizontal platforms approximately 6.5cm apart were recorded at 1000 fps using a highspeed camera (Fastcam SA3, Photron USA, Inc., San Diego, CA, USA) and a 105 mm 1:2.8 DG macro lens (Sigma Corp. of America, Ronkonkoma, NY, USA). Our analysis focused only on the takeoff phase, not the flight or landing phases of the jumps. The camera was oriented perpendicularly to the spider’s plane of motion, such that the right four legs were visible and in-focus throughout the jump. Videos in which any part of the right side of the spider moved out of the focal plane were discarded from analysis.

### Morphological measurements of spiders

After videos were recorded, photographs were taken of each animal; we were able to obtain live photographs of 42/46 spiders filmed (see Supplementary Figure S2). Following live photographs, spiders were sacrificed by submersion in 70% ethanol solution. Morphological measurements were taken from images of preserved spiders with legs separated from the body. Using FIJI [37], we traced a polygon around each of the following leg segments: femur, patella, tibia, metatarsus, and tarsus. Total leg lengths were calculated as the sum of the major axis of each leg segment polygon. Polygons were also traced around the cephalothorax and abdomen. Body length was calculated as the sum of the major axes for the polygons fit around the cephalothorax and abdomen. Any individuals for which morphological measurements were lacking were excluded from further analysis.

Individuals were identified to the genus level based on photographs of live individuals. Animals were not identified to the species level because preserved specimens were not available for the microscopic dissection and imaging that is necessary for this level of identification. A pruned phylogenetic tree (Figure 1) was generated based on the most recent published phylogeny of Salticidae [22].

### Video tracking

For body part tracking we used DeepLabCut 3 with the PyTorch engine [24, 31]. Specifically, we labeled 1141 frames taken from 28 videos/animals (of which 95% was used for training). We used a ResNet-50 based neural network with default parameters. We used three training iterations optimized each for 200 epochs with a batch size of 8. Data was incrementally increased from 436, 965, 1141 labeled frames where 95% was used for training and 5% for testing. At each iteration we used 1 shuffle. On the final iteration the test error was 29.29 pixels with train error of 18.4 pixels, corresponding to an error of approximately 1 mm. We then used a *p*-cutoff of 0.6 to condition the (*x, y*) coordinates for future analysis. This network was then used to analyze videos from similar experimental settings, including videos from previously published datasets on a North American species (*Habronattus conjunctus*) [6] and an Australian species (*Maratus rainbowi*) of jumping spider.

Final adjustments to DeepLabCut’s assigned points for all frames were carried out in a custom-designed GUI. A given joint label was adjusted if it met the following criteria: (1) the point was not located on the correct leg, (2) the point was not located within a 6-pixel-diameter circle of a human approximation of the joint location.

A total of 66 videos were analyzed across 46 individuals and 14 genera from the Amazonian dataset. For the later meta analysis (Figure 4), two videos each were used for the North American and Australian spiders.

A total of 25 points were tracked on each spider (Figure M1): 5 on the body (cephalothorax front, cephalothorax top, pedicel, abdomen top, abdomen back) and 5 along each leg (tip of the tarsus, tarsus-metatarsus joint, metatarsus-tibia joint, patella-femur joint and femur-trochanter joint). The tibia and patella are treated as a single element in our analysis as we did not see noticeable bending at the tibia-patella joint during takeoff.

**Figure M1:**
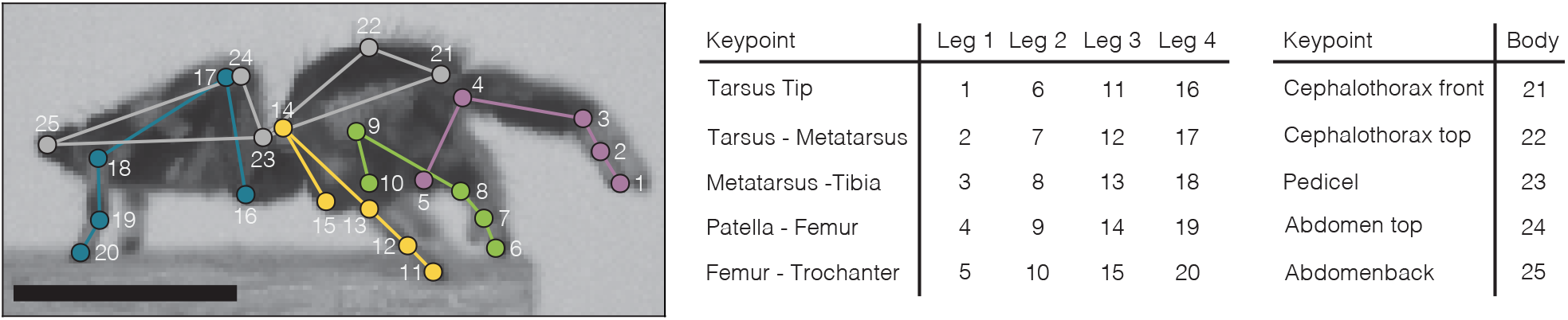
Tracked keypoints on the right side of the spider. The number of each point corresponds to the keypoint index in **x**_*i*_.

We denote the keypoint trajectories by **x**_*i*_(*t*) = (*x*_*i*_(*t*), *y*_*i*_(*t*)) where the mapping from point to index is given in M1. From the tracking we have each **x**_*i*_(*t*) sampled at the time points *t*_*i*_ = *i* ms.

### Kinematics analysis

#### Cropping the timeseries data

Each of the keypoint trajectories were cropped in time to center around the takeoff time. An approximate takeoff point was determined from the tracked tip of the third leg tarsus, **x**_11_. We approximate the speed of the point at each *t*_*i*_ by 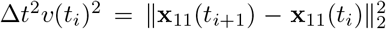, where Δ*t* is the constant time step between frames (1ms). The takeoff point is approximated as the transition point between *v*(*t*_*i*_) ≈0 and *v*(*t*_*i*_) *>* 0. To find the elbow where the transition happens we use a modified version of the kneedle algorithm [36]: the approximate velocity time series is normalized to lie between [0, 1] and smoothed using a median filter with window size 5 to give 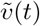. The transition point is chosen as the time point at the minimum of the difference curve 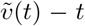. The time series are then cropped to keep 150ms before the takeoff point and then all frames after the jump. If a video did not have 150ms before the takeoff point then the full video is kept. The cropping prevents extended runs of the spider being stationary impacting downstream analysis.

#### Smoothing spline approximation of center of mass

The center of mass of the spider at each frame was approximated by the mean of the 5 body points at that frame, 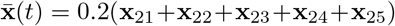. We approximate the spider’s velocity and acceleration by the time derivative of this center of mass 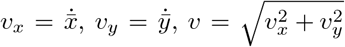 and 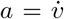. To compute the time derivatives we use smoothing cubic splines [34] modified to account for jitter. From the tracking we have the center of mass 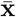 sampled at a set of uniform time points *t*_*i*_ = *i*Δ*t* for 0 ≤*i ≤ N*. It will be useful to stack the samples of each coordinates separately into vectors 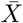 and 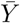, where 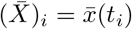 and similarly for 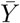. We approximate the continuous center of mass, **x**_cm_(*t*), using a polynomial spline of degree *D* of the form,

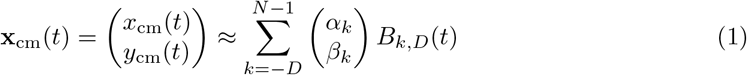

where *B*_*Dk*_(*t*) are the B-spline basis functions of degree *D* defined on the extended uniform knots 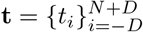 by the following recursion,

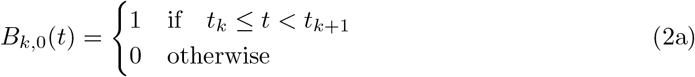

and

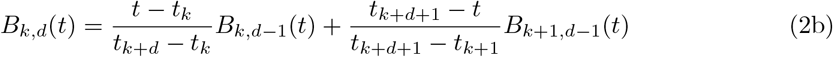

Here we choose to work with cubic *D* = 3 splines. To fit the spline to data we minimize the smoothing spline loss function separately for *x* and *y*,

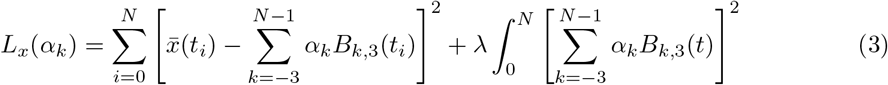

and similarly for *L*_*y*_(*β*_*k*_). We discuss the choice of *λ* in the next section.

#### Efficient fitting of smoothing spline

Applying this recursion on the uniform knots for *D* = 3 and using the natural boundary conditions, **x**^*′′*^(0), **x**^*′′*^(*N*) to extend the data to the extended nodes gives the B-spline matrix,

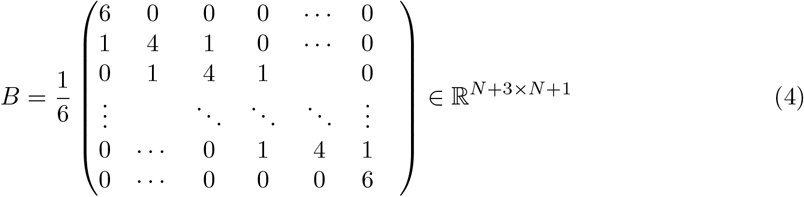

Using this matrix we can rewrite the loss functions as a regularized least squares optimization,

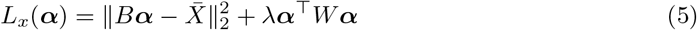

where *W* is constructed by integrating the pairwise combinations of B-splines and is given by

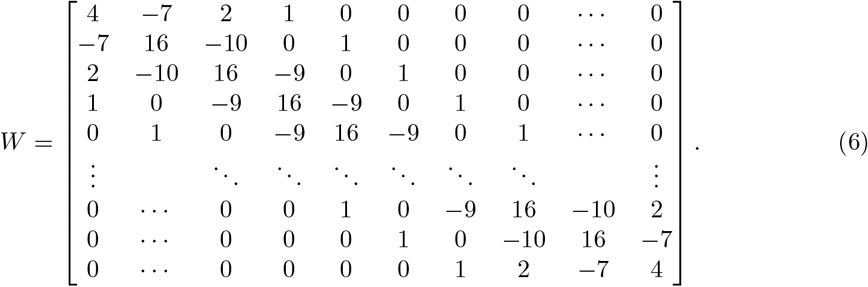

The solution of the least squares is given by the normal form,

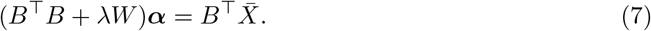

For a fixed value of *N* we can efficiently solve the least squares problem for multiple values of *λ* using the generalized eigen decomposition of *B*^⊤^*B* and *W*, such that *WV* = *B*^⊤^*BV D*, with *D* a diagonal matrix, to diagonalize the normal form of the equations [29]. Since *B*^⊤^*B* is symmetric positive definite *V* ^⊤^*B*^⊤^*BV* = *BV V* ^⊤^*B*^⊤^ = *I*. We define 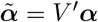 and 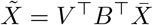, then the normal equations become

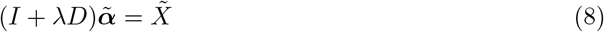

this can be solved for 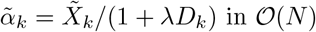.

#### Optimal value of regularization parameter

To determine the optimal value of *λ* we use the L-curve approach. In the tilde variables the curvature penalty is given by 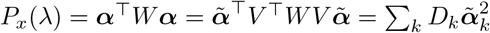. When we are trying to fit a smooth curve simultaneously we define a combined smoothness penalty,

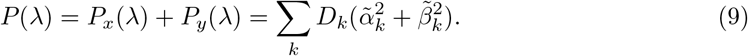

The residual term is

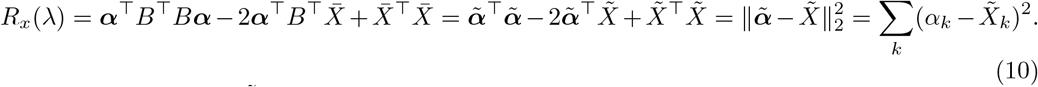

Substituting in 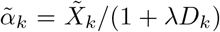 this reduces to 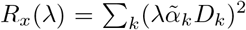. Again we define a combined residual,

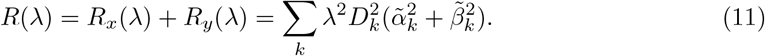

This means we can evaluate the two components of the loss function completely in the transformed tilde variable space. The L-curve is defined as the curve given by *x*(*λ*) = (1*/*2) log *R*(*λ*) and *y*(*λ*) = (1*/*2) log *P* (*λ*). We wish to find the elbow of this curve that balances the tradeoff between fitting the data well *x* small and the curve also being smooth *y* also small. As *λ* → 0, *x* → 0 and *y* will grow large, while as *λ* → ∞, *x* will grow large and *y* → 0. We define the optimal *λ* as the elbow of this curve given by its maximum curvature. We will make use of the following identity,

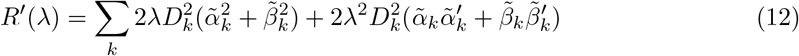

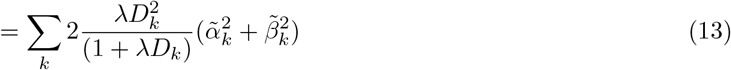

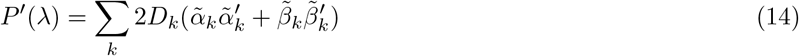

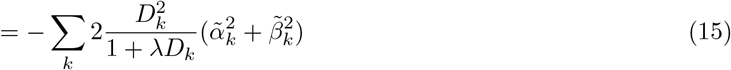

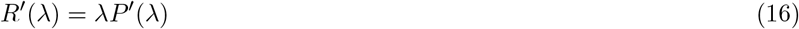

The curvature is given by *κ* = (*x*^*′*^*y*^*′′*^ −*x*^*′′*^*y*^*′*^)*/*(*x*^*′*2^ + *y*^*′*2^)^3*/*2^. Computing this with the identity above we have,

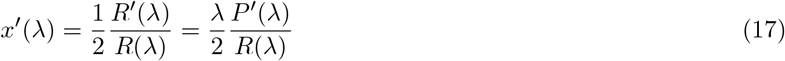

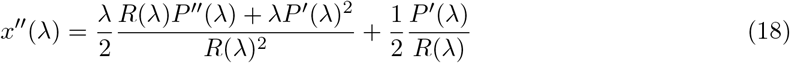

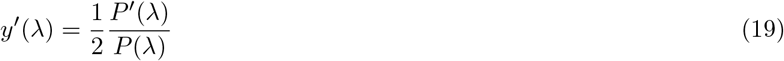

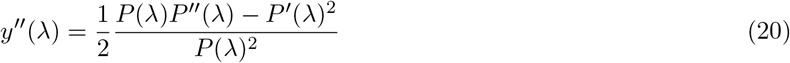

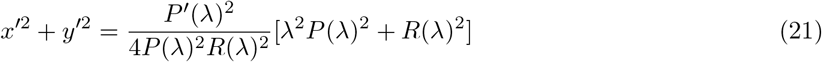

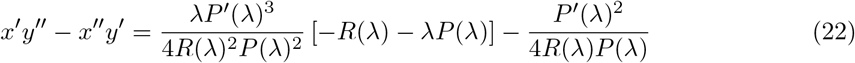

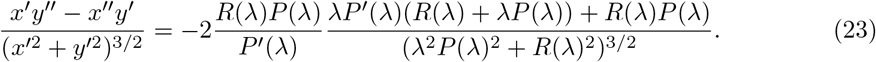

We can calculate *κ* efficiently using the values of *R, P*, and *P*^*′*^ that are also efficiently calculated directly in the tilde variables. This gives a fast *O*(*N*) method to calculate any point on the L-curve along with the corresponding curvature. The value of *λ* that maximizes the curvature can be calculated efficiently using classical one dimensional optimization techniques (see Figure M2 for an example L-curve and curvature). We use Brent’s method, implemented in Optim.jl, to minimize the cost function −*κ*(10^*l*^) over the range *l* ∈ [−4, 4]. Once the optimal *λ* is found the least squares problem is solved at this value of *λ* to find ***α, β***.

**Figure M2:**
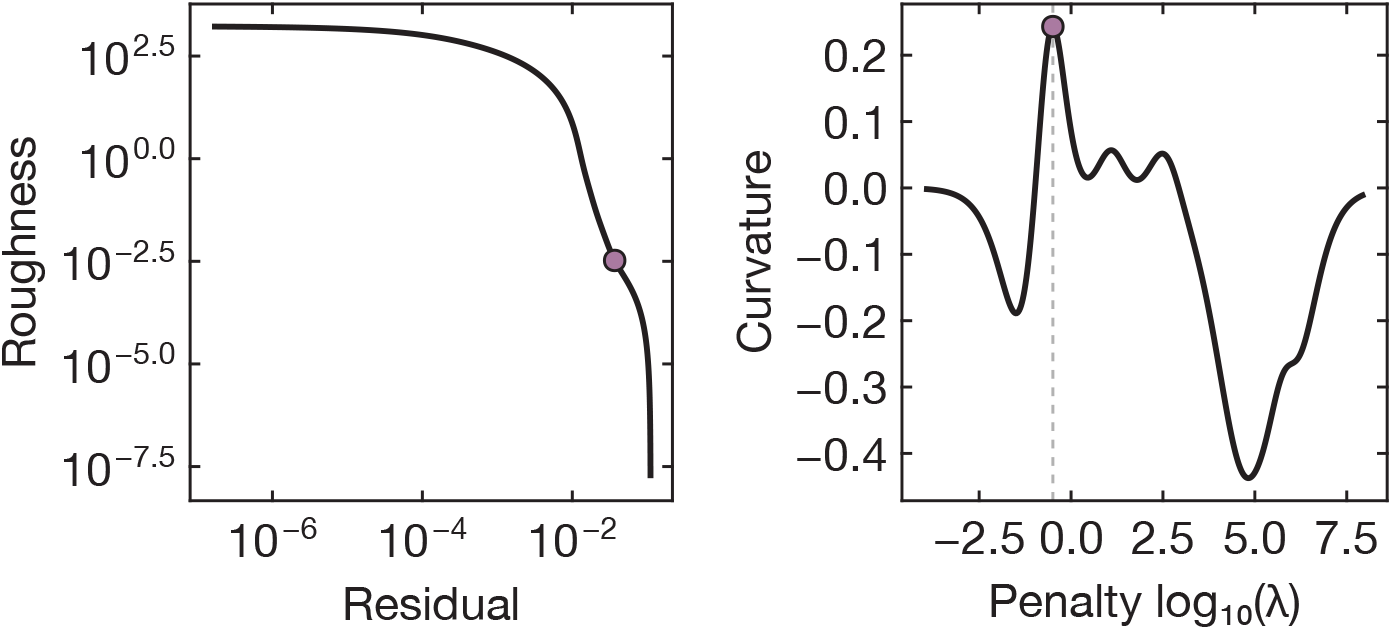
Example L-curve (left) shows a clear trade-off point. The L-curve curvature (right) has a distinct maximum value (purple circle) that corresponds to the elbow.

#### Jump detection and correction

A common occurrence in keypoint tracking is jitter, where at a given time point a systematic shift is introduced at time *t*_*i*_ and remains for all later times and outliers where a single time point is far from the trend. Such perturbations can significantly impact the quality of the cubic spline fit so we use a coupled approach to identify and correct for outliers and jitter.

To determine the positions of jumps in the trajectory we use the following procedure. First a smooth approximation is fit on the raw center of mass data to give ***α*** and ***β***. We then compute the residuals, 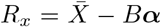 and 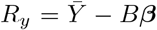. Since the smooth B-spline fit cannot rapidly adapt to jumps they also correspond to sudden jumps in the residual. To find this we define the difference magnitude 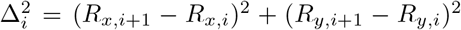. To determine is there is a jump at a given point we calculate the moving median 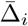 over a trailing window of length 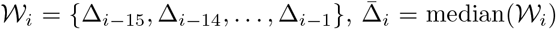 and also the median absolute deviation (mad),

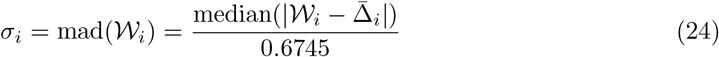

a robust measure of standard deviation, over the same. We say that there is a significant jump if the difference is greater that some number of *σ*_*i*_ from the baseline 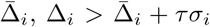. We chose a threshold of 15 to only remove clear and large jumps and outliers. We model the signal,

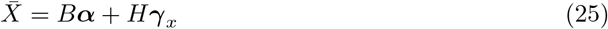

where *H* is a matrix constructed from Heaviside functions such that if the *j*th jump is identified at the *k*th time point, *H*_*i,j*_ = 0 for *i* ≤ *k* and *H*_*i,j*_ = 1 for *i > k*. The magnitude of the jump of *γ*_*j*_. To find the optimal jumps we solve the optimization problem,

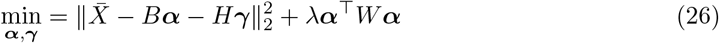

where we use the same *λ* as was used for the initial smoothing to compute the newest jump position. Setting the gradient with respect to ***α*** to 0 gives,

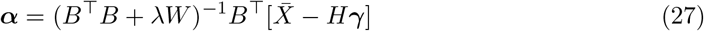

substituting this into the derivative with respect to ***γ*** set to 0 yields,

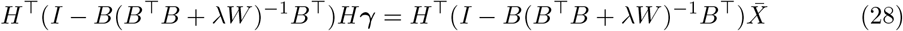

We notice that the term on the right hand side is exactly the residual *R*_*x*_ hence the the optimal jump can be found by solving the equation,

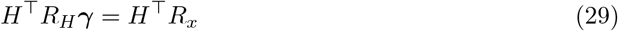

Every column of *R*_*H*_ is the residual of smoothing a step function at the same *λ* as used for *R*_*x*_ so can be computed efficiently using the previously described method. A equivalent procedure is used to fit the jump magnitude in *y*. Once we have found the optimal jumps ***γ*** we correct 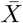 and 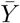 and rerun the smoothing spline procedure with a newly updated optimal *λ*. This procedure is repeated until no more jumps are detected. To find jumps in the initial window we then run the procedure on the time reversed signal and merge the jump locations. The final smooth approximation to the center of mass is given by the optimal smooth fit to the jump corrected data. See Figure M3 for an example of this procedure.

#### Derivative calculation

Once we have a fit to center of mass we compute the smooth velocities using, 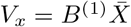 and 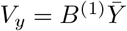 where *B*^(1)^ is the first derivative matrix of the B-spline matrix,

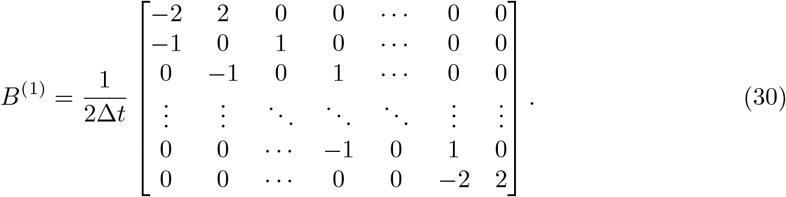

**Figure M3:**
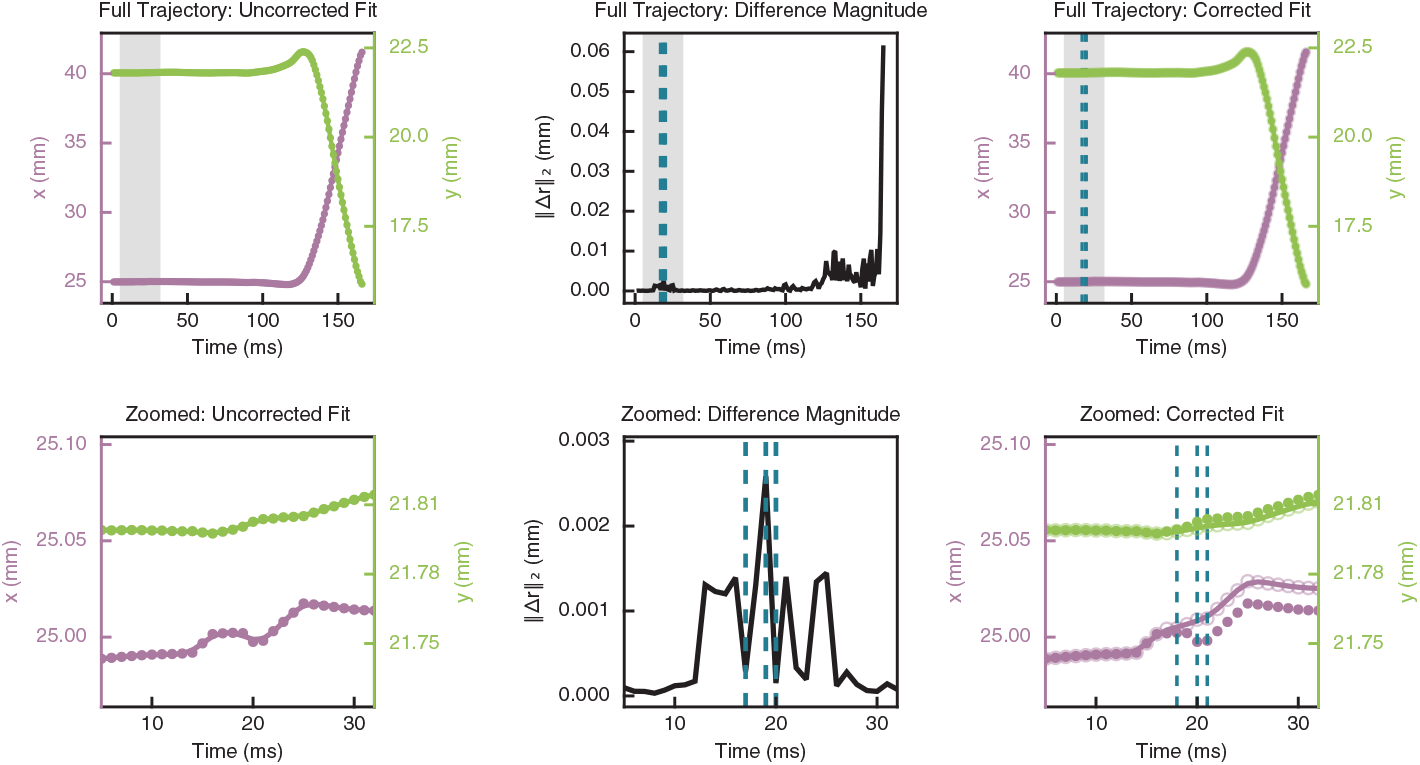
Example of the jump detection and correction. The left column is the raw center of mass trajectory (closed circle) with initial smoothing fit. The difference magnitude in the residual is shown middle column. Dashed lines represent positions where jumps are detected. Right column shows the corrected trajectory (open circles) and the updated fit. Bottom row is zoomed in (shaded gray region) for clarity.

From these velocity approximations we compute the speed vector *V* element-wise as 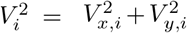. To compute the acceleration we compute the derivative of *V* by fitting a cubic spline to *V* but with a fixed small value of *λ* = 1.

#### Normalization of the velocity and acceleration data

Once we have the smooth cubic spline approximation, we nondimensionalize the kinematic measurement using the maximum value of the speed and by the jump duration. This gives a normalized curve where takeoff occurs at *t* = 0 and the jump start occurs at *t* = −1. We compute the jump duration by running a knee and elbow finding algorithm [36] on the speed vector *V*. We start by rescaling the velocity to lie between [0, 1], 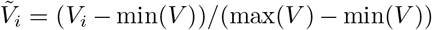. We then compute a difference curve where 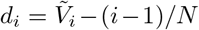 where *N* +1 is the length of vector *V*. We then define the start of the takeoff to be at the index where *d*_*i*_ is minimal *m* = argmin*d*_*i*_ and the takeoff speed to be at the index where *d*_*i*_ is maximal, *M* = argmax*d*_*i*_. To prevent issues at the boundary we enforce that *m < M*. The jump duration is then given by *T* = *t*_*M*_ −*t*_*m*_. We define a nondimensional time 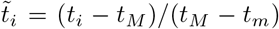 so that the jump occurs between 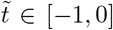. We nondimensionalize 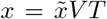 and 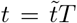. The nondimensional speed is then given by 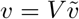 and the nondimensional acceleration is given by *a* = (*V/T*)*ã*.

#### Averaging kinematic trajectories

Every video now corresponds to nondimensionalized velocity and acceleration vectors sampled on slightly different normalized time grids 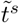. In order to average over these trajectories we apply a linear interpolation of the velocity and the acceleration to the same uniform time grid *t*_*u*_ ∈ [−1.8, 0.25]. No extrapolation is performed. When a video does not span the full range it is removed from the analysis of that time point. This only occurs after takeoff and we always have more than 55 out of the 66 samples at every time point. The median and quantiles are computed on the interpolated data excluding any missing points.

#### Effective leg length and joint angle

Effective leg length is computed by first approximating the length of the legs as

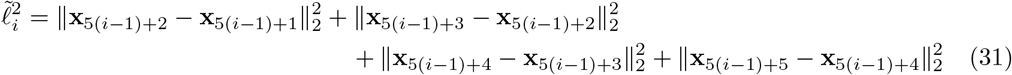

The effective leg length *e*_*i*_ is then given by the ratio of the distance from the tip of the leg to the end of the femur and the total leg length, 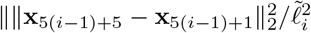. The length calculated from the keypoints was used to account for possible rotations relative to the plane of the camera.

The joint angle between the femur and patella was calculated using the keypoints by creating two vectors, the first r_*tp,i*_ = **x**_5(*i*−1)+3_ − **x**_5(*i*−1)+4_ that points along the tibia-patella away the joint and the other r_*f,i*_ = **x**_5(*i*−1)+5_ − **x**_5(*i*−1)+4_ that points along the femur away from the joint. We define two scalar quantities: the 2D cross product **r**_*f,i*_ × **r**_*tp,i*_ and the dot product **r**_*f,i*_ · **r**_*tp,i*_. We then calculate the angle 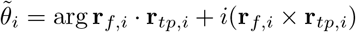, which is equivalent to atan2(**r**_*f,i*_ × **r**_*tp,i*_, **r**_*f,i*_ · **r**_*tp,i*_). This is a signed angle that is positive when the femur is in front of the patella. We finally define the angle as 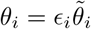 where *ϵ*_1_ = *ϵ*_2_ = −1 and *ϵ*_3_ = *ϵ*_4_ = 1 so that *θ*_*i*_ is positive when the leg is extended and negative when the joint is flexed.

Quantiles and medians for effective leg length and joint angle are computed as for the velocity and acceleration.

#### Choice of jumping regions

We break the graph into 5 distinct domains in normalized time to guide the comparisons. The transitions between these points is chosen as as follows 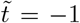 (start of take off), 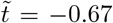 (peak in leg 2 effective leg length when front leg extension is maximum), 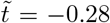 (elbow of leg 4 effective leg length when leg 4 is approximately fully extended), 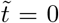 (the approximate point of take off).

#### Morphological and kinematic comparisons

For morphological and kinematic comparisons leg lengths from the segmented spiders as described in Morphological measurements of spiders. The jump duration and maximum speed are the same as used for nondimensionalization of the trajectories.

### Graph based modal analysis of keypoints

To further analyze the inter-limb coordination of the spiders during jumping we employ a modal analysis of the keypoints based on the graph shown in 3(A, right). The nodes correspond to the tracked keypoints and the connections to anatomical connections. We will use *n*_*i*_ to refer to the node corresponding to the keypoint position **x**_*i*_ and *e*_*ij*_ to refer to the edges between nodes *n*_*i*_ and *n*_*j*_. We define the position functions on the graph as *X*(*n*_*i*_, *t*) = *x*_*i*_(*t*) and *Y* (*n*_*i*_, *t*) = *y*_*i*_(*t*). On the graph we use the standard inner product. Given two functions *f* and *g* defined on the graph their inner-product ⟨*f, g*⟩ = _*i*_ *f* (*n*_*i*_)*g*(*n*_*i*_). On the graph we define the compactly supported orthonormal basis functions *ϕ*_*m*_(*n*_*i*_). Our basis is inspired by the Haar wavelets adapted to the spider graph. The basis is constant on every non zero section and iteratively focuses on more localized parts of the graph with compact support. We define the vectors ***ϕ***_*i*_ such that the *j*th element is *ϕ*_*i*_(*n*_*j*_). We can roughly break this basis functions into groups. The first group gives the interlimb and body coordination:

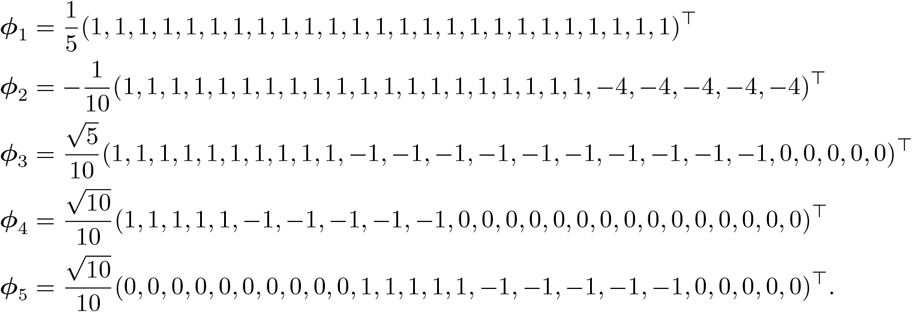

The second group gives information about the body: we will use **0**_*N*_ to represent a block of *N* repeated zeros,

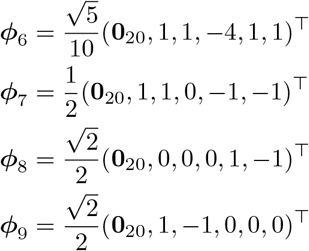

The final blocks are along each leg. For each leg block *k* ∈ {0, 1, 2, 3} the same pattern is shifted:

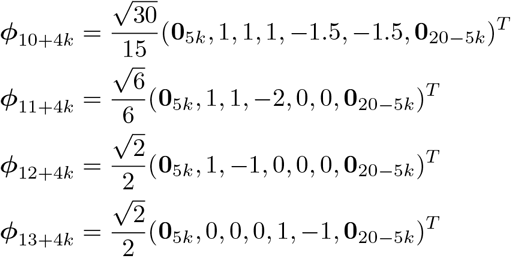

See Figure M4 for the complete basis on the spider graph.

We stack all the *x* coordinates at a given time point into the vector **X** such that *X*_*i*_(*t*) = *x*_*i*_(*t*) and similarly for *y* into **Y**, *Y*_*i*_(*t*) = *y*_*i*_(*t*). We can then calculate the wavelet coefficients corresponding to a given mode by 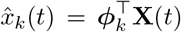 and 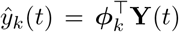. Since the basis is orthonormal and trajectory can be recreated at,

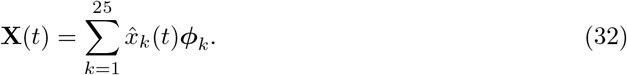

From this picture we see that all of the temporal information about the keypoints is pushed into the coefficient trajectories while information about the relative arrangement of the keypoints is determined by the basis modes.

#### Normalization of modal coefficients

These coefficients are not rotationally invariant, so we work instead with the rotationally invariant measure 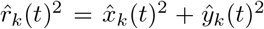. The wavelet coefficients were again interpolated to uniform time using linear interpolation and normalized by subtracting the median in the time normalized time window [−1.8, 0.25] and then dividing by the interquantile range between the 95th and 5th quantiles. From these scaled coefficients the median and bands are calculated as for the kinematic measurements.

**Figure M4:**
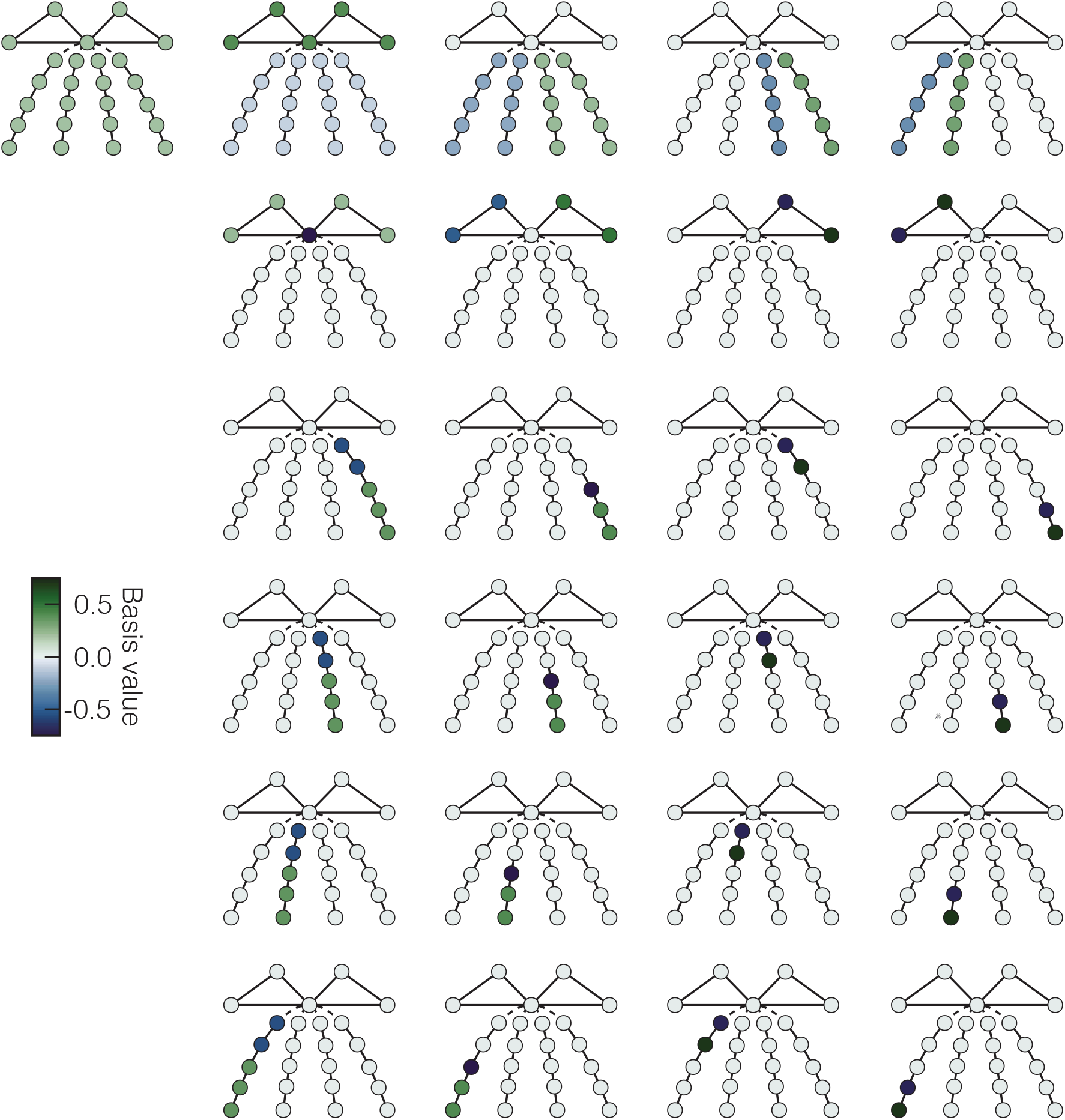
Full wavelet basis for the spider graph.

#### Other choices of basis function

We did not use the graph Laplacian basis, a standard orthogonal basis on graphs as this does not have compact support and as such is less interpretable in the biomechanical context than the constructed Haar inspired graph. See Figure M5 for the graph Laplacian basis computed on the

**Figure M5:**
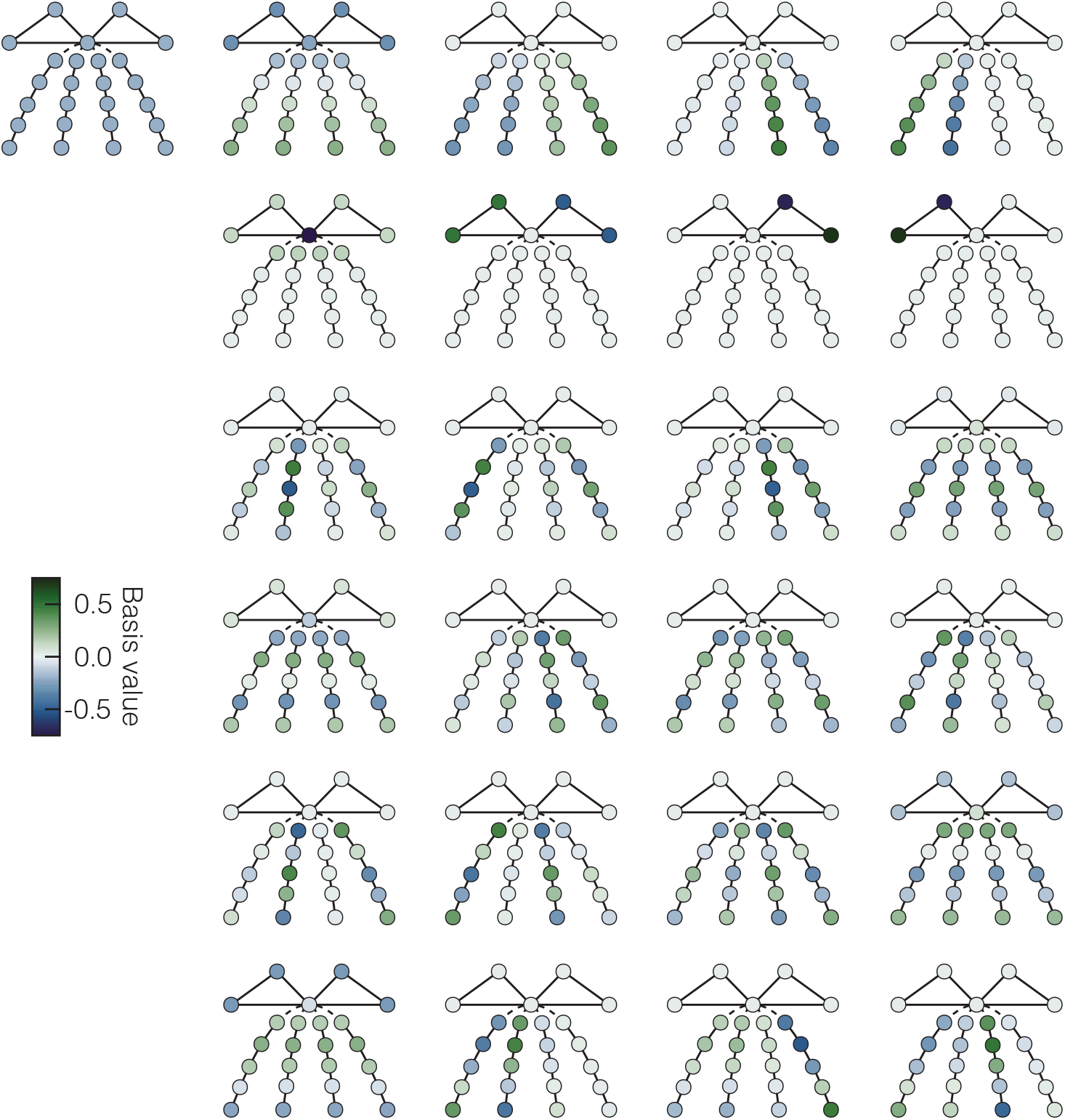
Full graph Laplacian basis for the spider graph does not have compact support like the wavelet basis.

### Analysis of published data

The data from the North American and Australian spiders were analyzed in the same way as for the Peruvian spiders for the plots in Fig. 4.

## Data Availability

All data and code in this paper will be made available upon publication.

## Acknowledgments

E.E.B., A.D.H., and J.A.N. acknowledge funding and support by NSF-Simons National Institute for Theory and Mathematics in Biology, which is jointly supported by the U.S. National Science Foundation (Award #2235451) and the Simons Foundation (Award MP-TMPS-00005320). J.A.N. and E.E.B. acknowledge support from the National Science Foundation through the Center for Living Systems (Grant #2317138). Data in this paper were collected by L.Y., J.H., L.B.A., and M.S.B at the Jungle Biomechanics Lab funded by NSF-IRES #2246236. We also gratefully acknowledge Gustavo Ruiz, who performed most of the species identification work, and Mariya Savinov, who provided helpful feedback on early versions of the analysis in this study.

