## Supplementary Figures 1-5 for "Takeoff dynamics are stereotyped across jumping spiders"

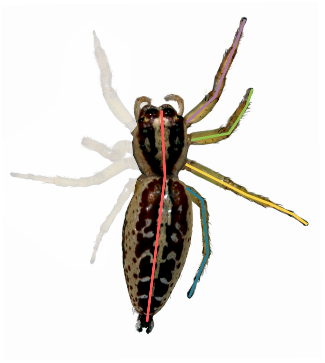

$$\frac{l1}{l1+l2+l3+l4} = \text{relative leg length } (l1)$$

$$\frac{bl}{\left(\frac{l1+l2+l3+l4}{4}\right)} = \text{legginess}$$

Figure S1: Calculation of relative leg length and legginess. Relative leg length is calculated as the length of a leg divided by the sum of all legs. Legginess is calculated as the body length (cephalothorax length + abdomen length) divided by mean leg length. Contralateral pairs of legs are assumed to be identical in length. Each leg length is the mean of both left and right leg in contralateral pair.

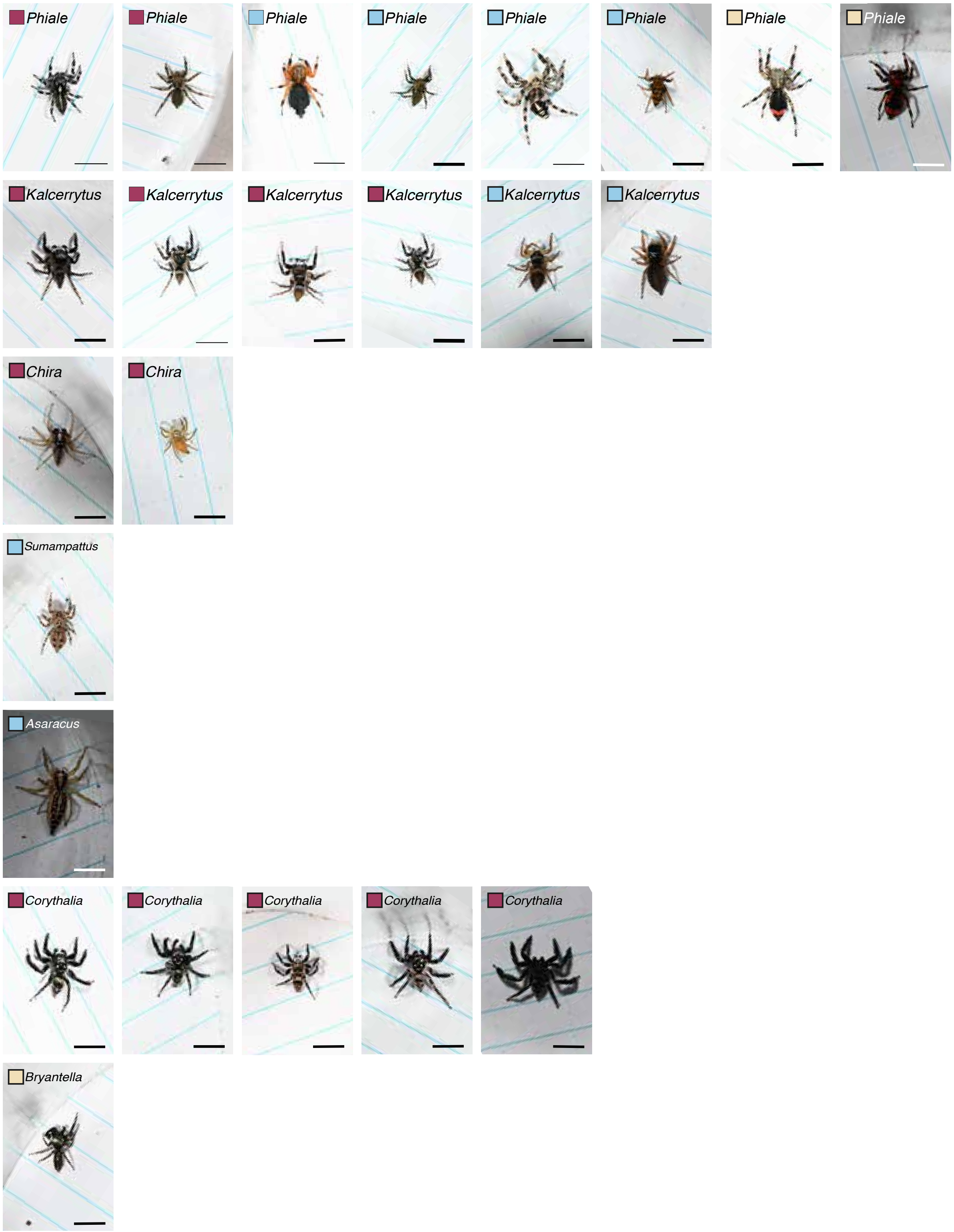


Figure S2a: Live images of 42/46 individuals used to construct the phylogeny and morphology measurements in Figure 1. Maroon box indicates mature male individual, light blue indicates mature female, and pale yellow indicates undetermined sex and/or immature. Figure continues next page.

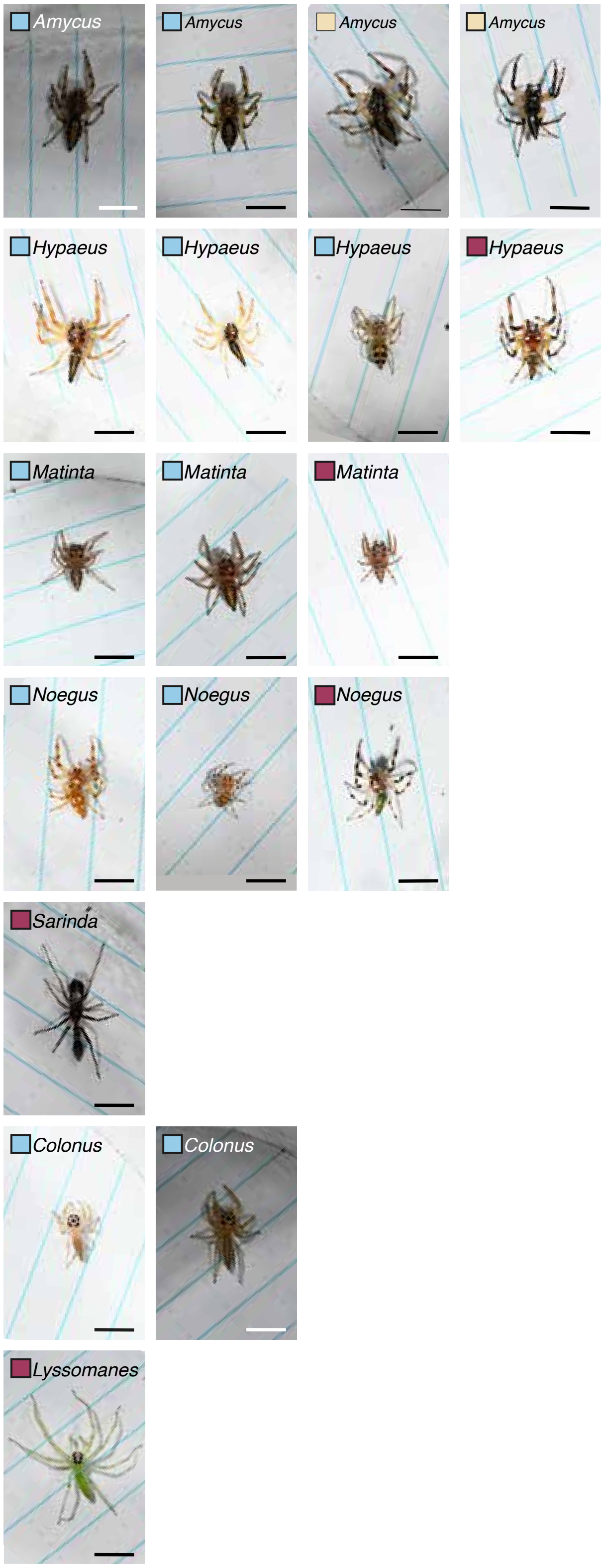


Figure S2b: Live images of 42/46 individuals used to construct the phylogeny and morphology measurements in Figure 1. Maroon box indicates mature male individual, light blue indicates mature female, and pale yellow indicates undetermined sex and/or immature.

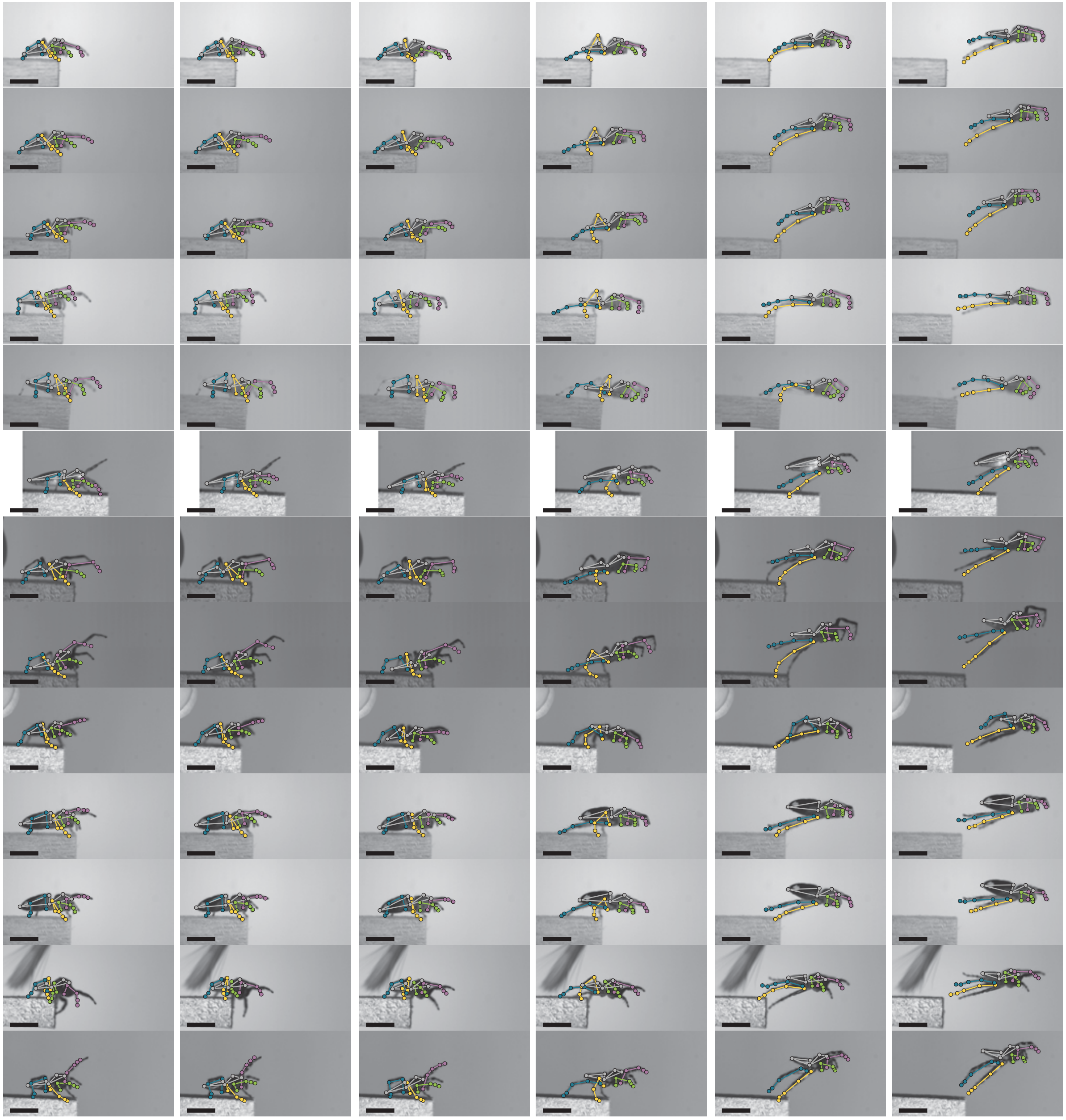

Figure S3a: All 66 videos used in the analysis aligned to the dashed lines positions in Figure 2. Last panel is the last time point in the video. Figure continued on next pages (1/5).

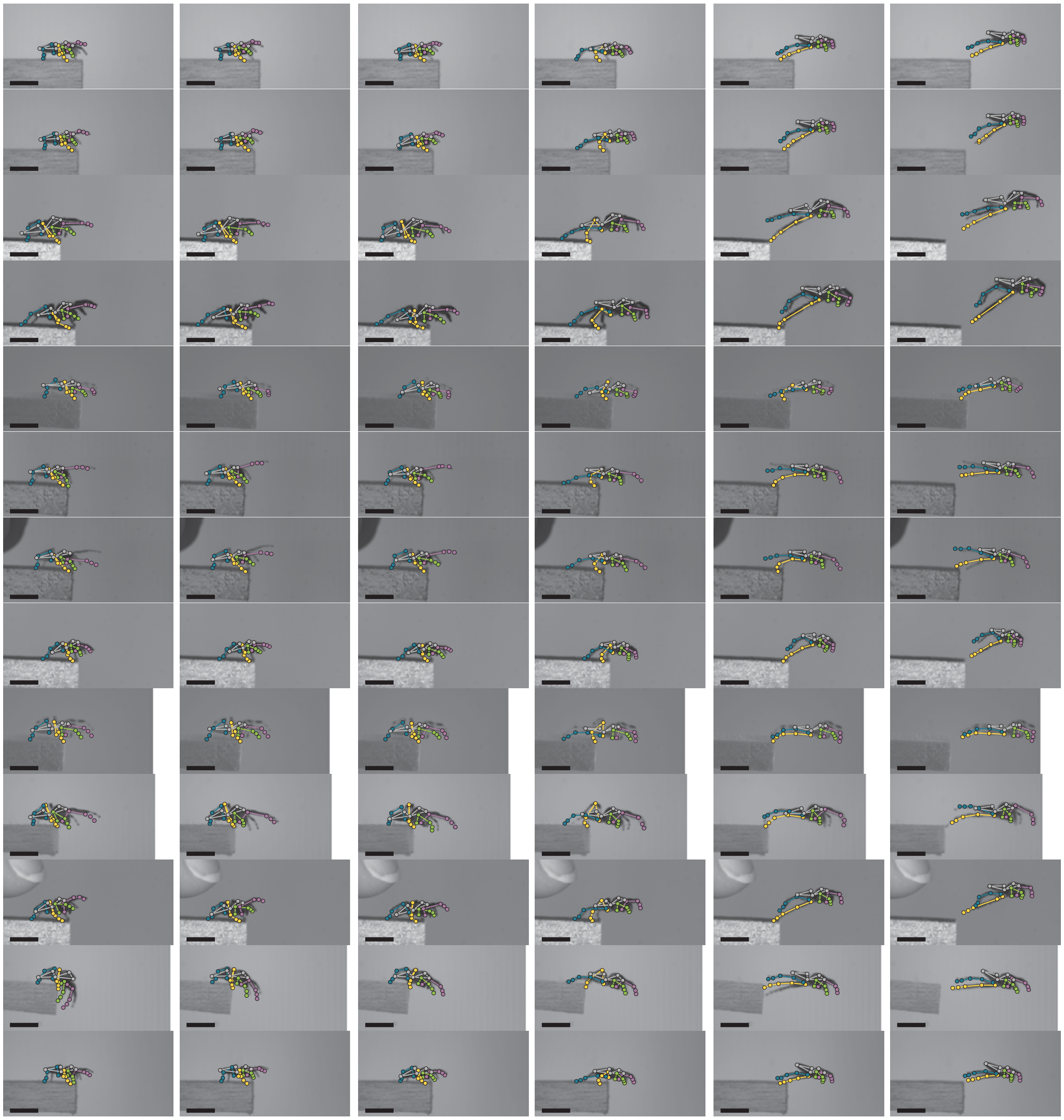

Figure S3b: All 66 videos used in the analysis aligned to the dashed lines positions in Figure 2. Last panel is the last time point in the video. Figure continued on next pages (2/5).

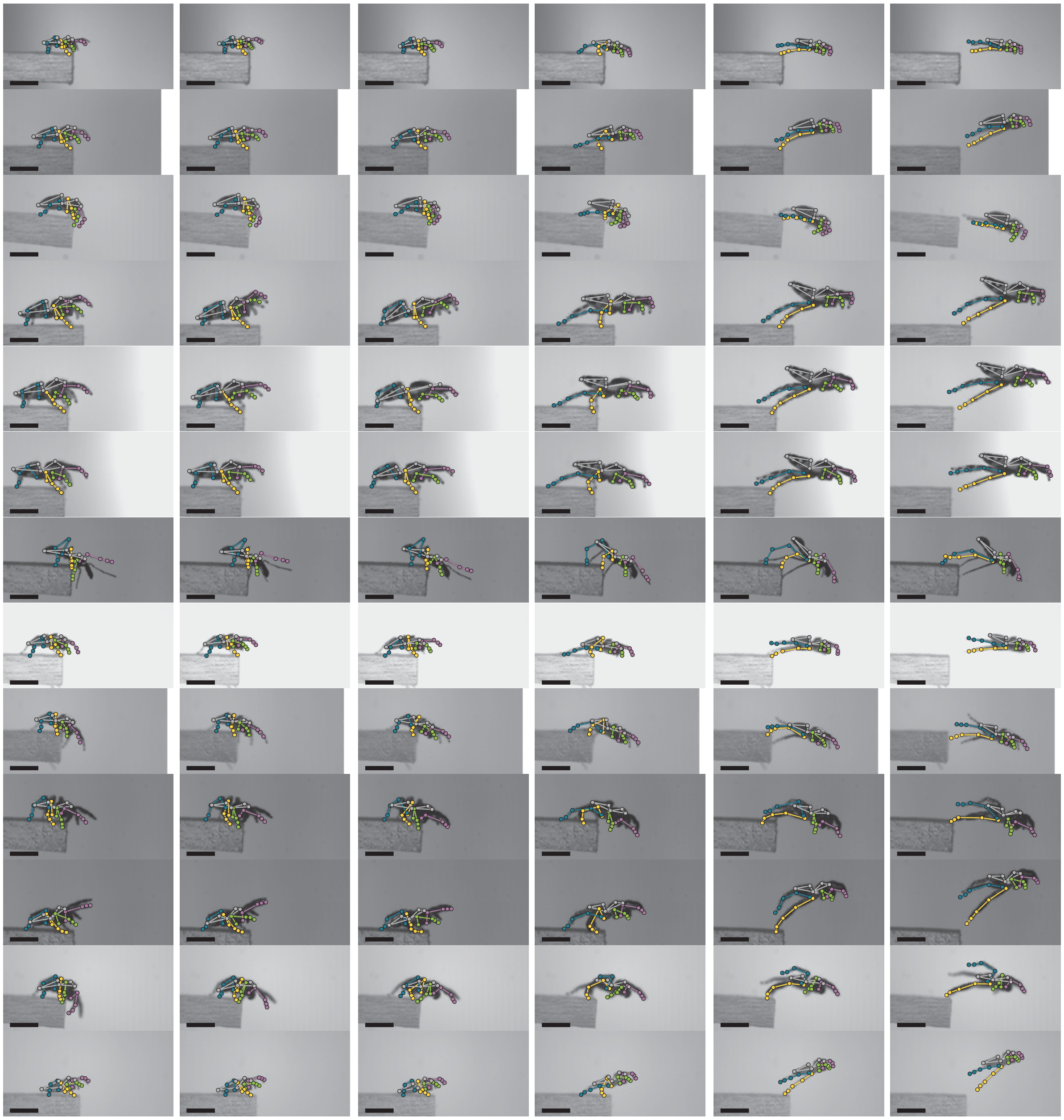

Figure S3c: All 66 videos used in the analysis aligned to the dashed lines positions in Figure 2. Last panel is the last time point in the video. Figure continued on next pages (3/5).

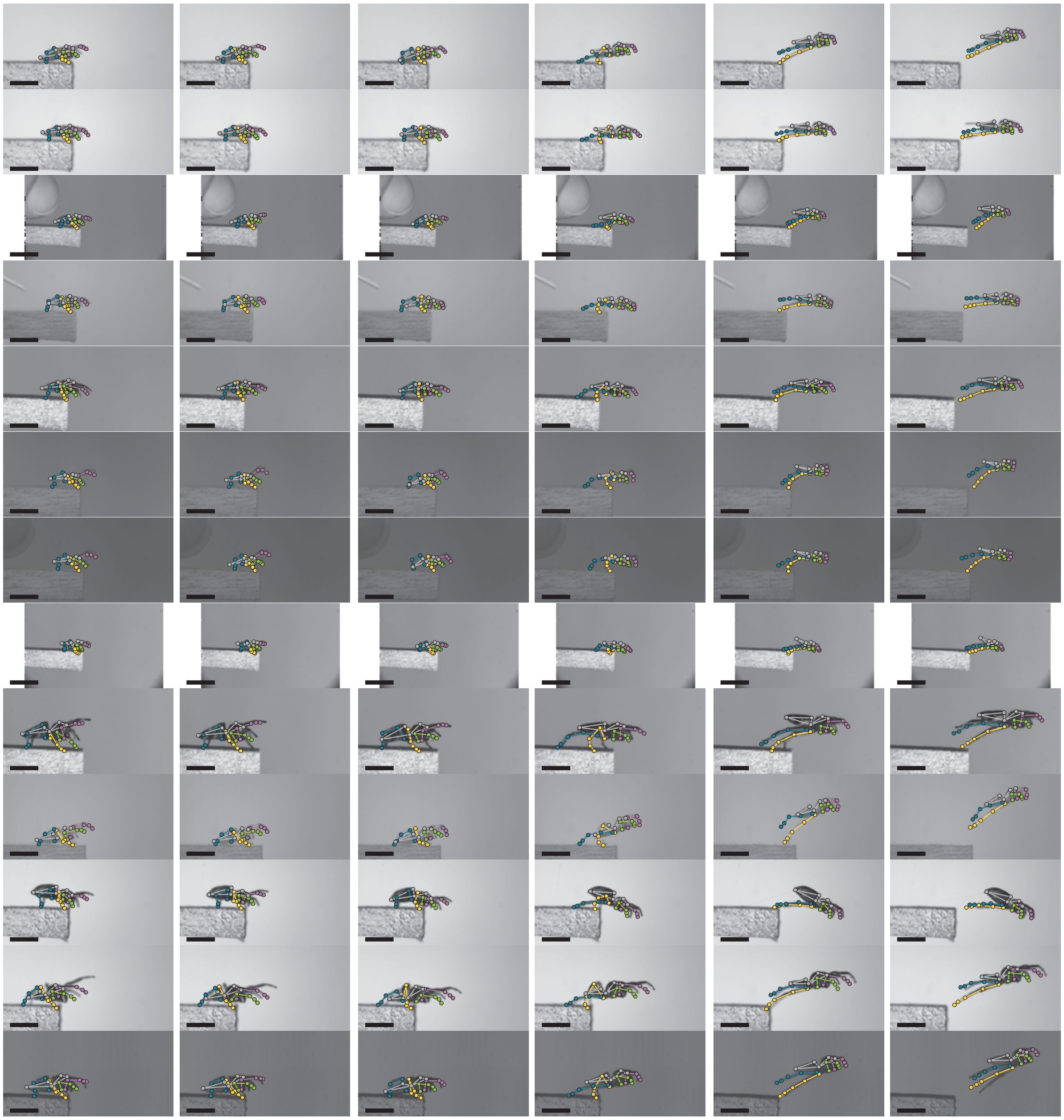

Figure S3d: All 66 videos used in the analysis aligned to the dashed lines positions in Figure 2. Last panel is the last time point in the video. Figure continued on next pages (4/5).

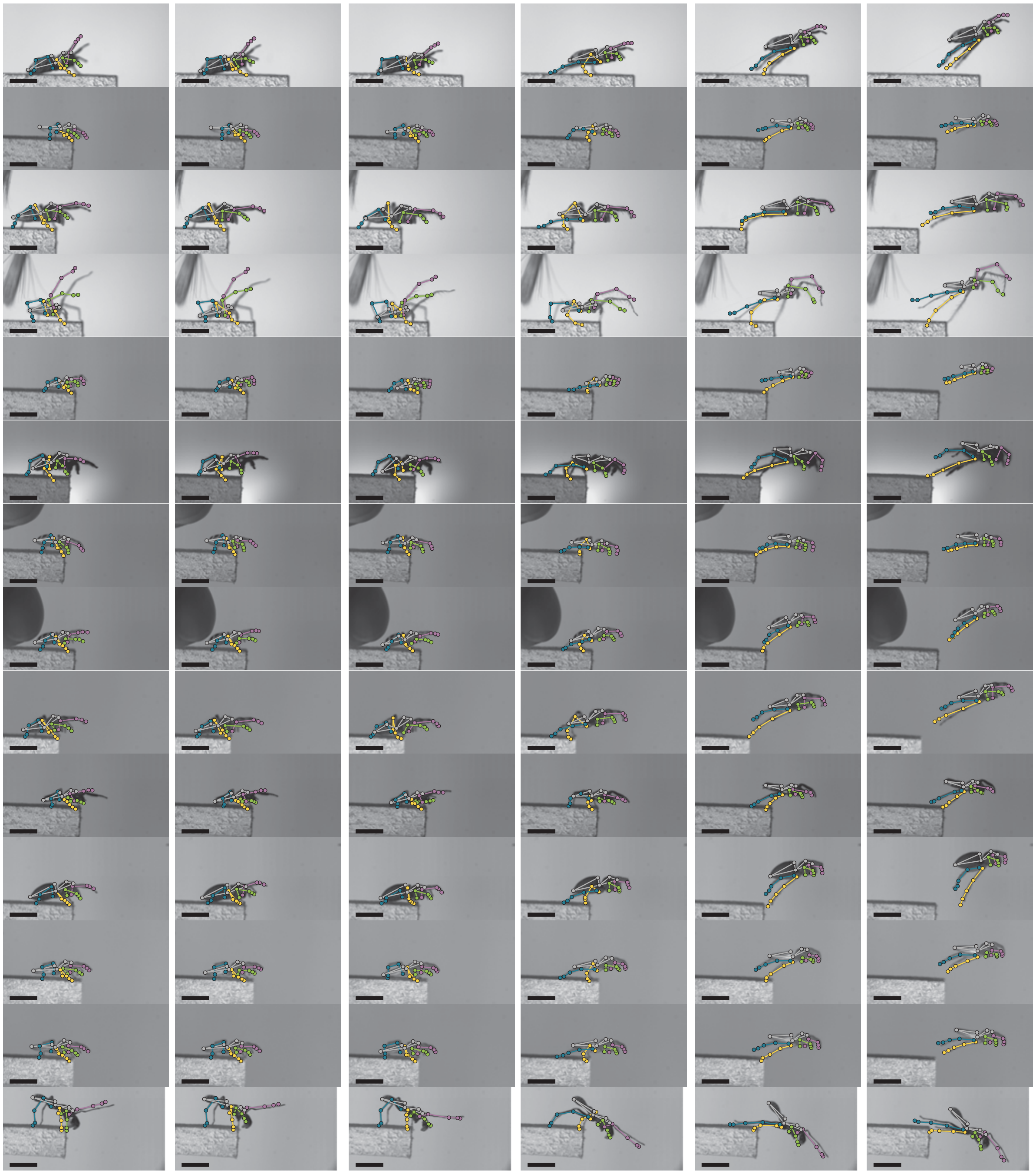

Figure S3e: All 66 videos used in the analysis aligned to the dashed lines positions in Figure 2. Last panel is the last time point in the video. (5/5)

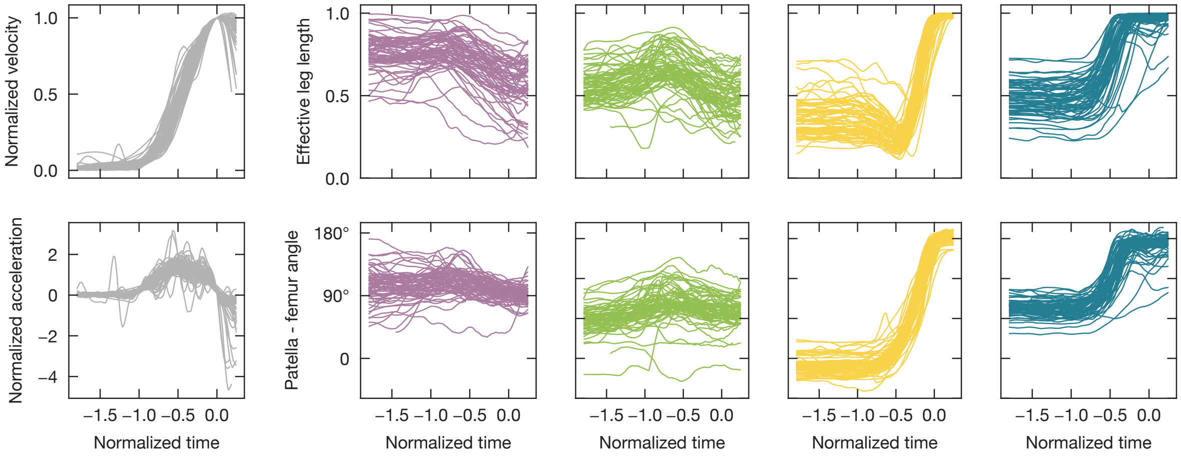


Figure S4: Normalized kinematic trajectories corresponding to the quantile bands shown in Figure 2. Each line corresponds to a separate video trial.

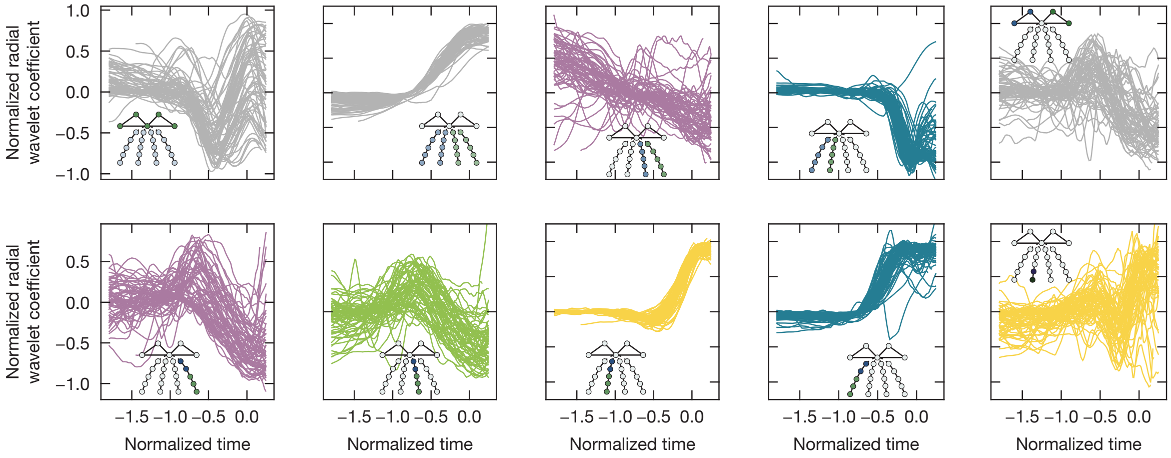


Figure S5: Normalized wavelet coefficient trajectories corresponding to the quantile bands shown in Figure 3. Each line corresponds to a separate video trial.
